# Selective maintenance of multiple CRISPR arrays across prokaryotes

**DOI:** 10.1101/148544

**Authors:** Jake L. Weissman, William F. Fagan, Philip L.F. Johnson

## Abstract

Prokaryotes are under nearly constant attack by viral pathogens. To protect against this threat of infection, bacteria and archaea have evolved a wide array of defense mechanisms, singly and in combination. While immune diversity in a single organism likely reduces the chance of pathogen evolutionary escape, it remains puzzling why many prokaryotes also have multiple, seemingly redundant, copies of the same type of immune system. Here, we focus on the highly flexible CRISPR adaptive immune system, which is present in multiple copies in a surprising 28% of the prokaryotic genomes in RefSeq. We use a comparative genomics approach looking across all prokaryotes to demonstrate that, on average, organisms are under selection to maintain more than one CRISPR array. We hypothesize that a tradeoff between memory span and learning speed could select for both “long-term memory” and “short-term memory” CRISPR arrays, and we go on to develop a mathematical model to show that such a tradeoff could, in theory, lead to selection for multiple arrays.

## Introduction

Just as larger organisms must cope with the constant threat of infection by pathogens, so too must bacteria and archaea. To defend themselves in a given pathogenic environment, prokaryotes may employ a range of different defense mechanisms, and oftentimes more than one (Makarova et al., 2011b, 2013; Houte et al., 2016). While having multiple types of immune systems may decrease the chance of pathogen evolutionary escape (Iranzo et al., 2015), having multiple instances of the same type of system is rather more puzzling. Here we explore this apparent redundancy in the context of CRISPR-Cas immunity.

The CRISPR-Cas immune system is a powerful defense mechanism against the viruses that infect bacteria and archaea and is the only known example of adaptive immunity in prokaryotes (Makarova et al., 2006; Goren et al., 2012). This system allows prokaryotes to acquire specific immune memories, called “spacers”, in the form of short viral genomic sequences which they store in CRISPR arrays in their own genomes (Mojica et al., 2005; Bolotin et al., 2005; Barrangou et al., 2007). These sequences are then transcribed and processed into short crRNA fragments that guide CRISPR-associated (Cas) proteins to the target viral sequences (or “protospacers”) so that the foreign DNA or RNA can be degraded (Barrangou et al., 2007; Marraffini and Sontheimer, 2008; Marraffini, 2015). Thus the CRISPR array is the genomic location in which memories are recorded, while the Cas proteins act as the machinery of the immune system, with specific proteins implicated in memory acquisition, crRNA processing, or immune targeting.

CRISPR systems appear to be widespread across diverse bacterial and archaeal lineages, with previous analyses of genomic databases indicating that ∼40% of bacteria and ∼80% of archaea have at least one CRISPR system (Makarova et al., 2011a; Sorek et al., 2013; Burstein et al., 2017). These systems vary widely in *cas* gene content and targeting mechanism, although the *cas1* and *cas2* genes involved in spacer acquisition are universally required for a system to be fully functional (Barrangou et al., 2007; Makarova et al., 2011a). Such prevalence suggests that CRISPR systems effectively defend against phage in a broad array of environments. The complete story seems to be more complicated, with recent analyses of environmental samples revealing that some major bacterial lineages almost completely lack CRISPR systems and that the distribution of CRISPR systems across prokaryotic lineages is highly uneven (Burstein et al., 2016). Other studies suggest that particular environmental factors can be important in determining whether or not CRISPR immunity is effective (e.g., in thermophilic environments Iranzo et al. 2013; Weinberger et al. 2012b). While previous work has focused on the presence or absence of CRISPR across lineages and habitats, little attention has been paid to the number of systems in a genome.

In fact, the multiplicity of CRISPR systems per individual genome varies greatly, with many bacteria having multiple CRISPR arrays and some having multiple sets of *cas* genes as well (e.g., Horvath et al. 2009; Cai et al. 2013). CRISPR and other immune systems are horizontally transferred at a high rate relative to other genes in bacteria (Puigbò et al., 2017), meaning that any apparent redundancy of systems may simply be the result of the selectively neutral accumulation of systems within a genome. Alternatively, there are a number of reasons, discussed below, why having multiple sets of *cas* genes or CRISPR arrays might be under selection.

We suspected that prokaryotes may be under selection to maintain multiple CRISPR arrays, given that it is common for organisms across lineages to have multiple systems (as detailed below) and, in some clades, these appear to be conserved over evolutionary time (e.g. Boudry et al. 2015; Andersen et al. 2016). Because microbial genomes have a deletion bias (Mira et al., 2001; Kuo and Ochman, 2009), we would expect extraneous systems to be removed over time. Here we construct a test of neutral CRISPR array accumulation via horizontal transfer and loss. Using publicly available genome data we show that the number of CRISPR arrays in a wide range of prokaryotic lineages deviates from this neutral expectation by approximately two arrays. Thus we conclude that, on average, prokaryotes are under selection to have multiple CRISPR arrays. We go on to discuss several hypotheses for why having multiple arrays might be adaptive. Finally, we suggest that a tradeoff between the rate of acquisition of immune memory and the span of immune memory could lead to selection for multiple CRISPR arrays.

## Materials and methods

### Dataset

All available completely sequenced prokaryotic genomes (all assembly levels) were downloaded from NCBI’s non-redundant RefSeq database FTP site (ftp://ftp.ncbi.nlm.nih.gov/genomes/refseq/bacteria, O’Leary et al. 2016) on December 23, 2017. Genomes were scanned for the presence of CRISPR arrays using the CRISPRDetect software (Biswas et al., 2016). We used default settings except that we did not take the presence of *cas* genes into account in the scoring algorithm (to avoid circularity in our arguments), and accordingly used a quality score cutoff of three, following the recommendations in the CRISPRDetect documentation. CRISPRDetect also identifies the consensus repeat sequence and determines the number of repeats for each array. Presence or absence of *cas* genes were determined using genome annotations from NCBI’s automated genome annotation pipeline for prokaryotic genomes (Tatusova et al., 2016). We discarded genomes that lacked a CRISPR array in any known members of their taxon. In this way we only examined genomes known to be compatible with CRISPR immunity.

### Test for selection maintaining multiple arrays

Our power to detect selection hinges on our ability to differentiate between non-functional (i.e., neutrally-evolving) and functional (i.e., potentially-selected) CRISPR arrays. Since all known CRISPR systems require the presence of *cas1* and *cas2* genes in order to acquire new spacers, we use the presence of both genes as a marker for functionality and the absence of one or both genes as a marker for non-functionality. Henceforth we will consider CRISPR arrays in genomes with both *cas1* and *cas2* genes to be “functional” and CRISPR arrays in genomes lacking either *cas1, cas2*, or both genes to be “non-functional”. This differentiation allows us to consider the probability distributions of the number of CRISPR arrays *i* in non-functional (*N*_*i*_) and functional (*F*_*i*_) genomes, respectively.

We start with our null hypothesis that in genomes with functional CRISPR systems possession of a single array is highly adaptive (i.e. viruses are present and will kill any susceptible host) but additional arrays provide no additional advantage. Thus these additional arrays will appear and disappear in a genome as the result of a neutral birth/death horizontal transfer and loss process, where losses are assumed to remove an array in its entirety. This hypothesis predicts that the non-functional distribution will look like the functional distribution shifted by one (*S*_*i*_):

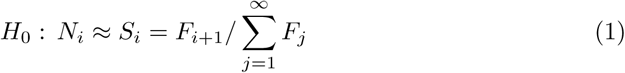

for *i* ≥ 0. We take two approaches to testing this prediction: one parametric from first principles with greater power but more assumptions and one non-parametric with less power but also fewer assumptions.

We begin by deriving a functional form for the distribution *N*_*i*_ from first principles following a neutral process. If CRISPR arrays arrive in a given genome at a constant rate via rare horizontal transfer events, then we can model their arrivals using a Poisson process with rate *η*. Assuming arrays are also lost independently at a constant rate, the lifetime of each array in the genome will be exponentially distributed with rate *ν*. This leads to a linear birth-death process of array accumulation, which yields a Poisson equilibrium distribution with rate 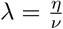. While this rate might be constant for a given taxon, it will certainly vary across taxa due to different intrinsic (e.g. cell wall and membrane structure) and extrinsic factors (e.g. density of neighbors, environmental pH and temperature) (Puigbò et al., 2017). We model this variation by allowing genome *j* to have rate 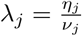 and assuming *λ*_*j*_ ∼ Gamma(*α, β*), which we pick for its flexibility and analytic tractability. This combination of gamma and Poisson distributions leads to the number of arrays *i* in a random genome following a negative binomial distribution *N*_*i*_ = NB(*r, p*) where *r* = *α* and 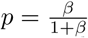.

Now we can fit this distribution to data to find maximum likelihood estimates of *r* and *p* for the distribution of array counts in both the set of non-functional genomes (*N*_*i*_) and the set of functional genomes as shifted under our null hypothesis (*S*_*i*_). This allows us to construct a parametric test of multi-array adaptiveness. We expect that 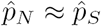 and 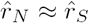 under our null hypothesis (where subscripts correspond to the distribution to which the parameters were fit). When our null hypothesis is violated it should shift the means of these distributions. Therefore we estimate and compare these means 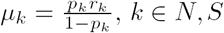. We expect that 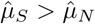 if more than one array is selectively maintained, and we bootstrap confidence intervals on these estimates to determine whether the effect is significant.

We also construct a non-parametric test for selection by determining at what shift *s* the mismatch between 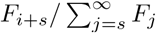 and *N*_*i*_, measured as the sum of squared differences between the distributions, is minimized:

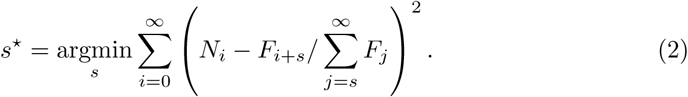

Under our null hypothesis *s*\* = 1, and a value of *s*\* > 1 implies that selection maintains more than one array. Our parametric test is superior to *s*\* because it can detect if selection maintains more than one array across the population on average, but not in all taxa, so that the optimal shift is fractional.

### Correcting for phylogenetic correlations in HGT

Differential rates of horizontal gene transfer (HGT) between lineages could produce an observed correlation between *cas1* and *cas2* presence and array count in the absence of any selection for having multiple CRISPR arrays. In other words, some genomes would be functional and have many arrays due to a high arrival rate of foreign genetic material, and other lineages would be non-functional and lack CRISPR arrays simply because of low rates of HGT. If this were the case, then comparisons between these lineages would lead to a spurious result of selection.

There are several ways to control for such correlation. First, we can perform our parametric test on a subset of the data such that we take an equal number of functional and non-functional genomes from each species to control for lineage-specific effects. Second, we can also perform a species-wise parametric test. In this case, for each species *k* we calculate 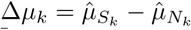 and then bootstrap the mean of the distribution of these values 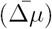 to detect if there is a significant difference from zero. We also map these values onto a phylogeny (SILVA Living Tree 16s rRNA tree; Yarza et al. 2008) and perform a formal test for phylogenetic signal (the *K* statistic; Blomberg et al. 2003; Revell 2012), which indicates whether or not any signal of selection is isolated to a particular portion of the tree.

Finally, our test for selection can also be conceptualized of in terms of a regression, which has standard methods for phylogenetic correction. Essentially, if we fit a model of array counts per genome as predicted by functionality, our null hypothesis states that the slope of the regression should be approximately one, with a slope larger than one indicating that selection maintains multiple arrays. We fit a linear model of mean number of arrays in a genome per species predicted by the proportion of functional genomes per species. We then fit a linear model accounting for phylogeny (using the phylolm R package and assuming a Brownian motion model for trait evolution; Tung Ho and Ané 2014), again using the SILVA tree and limiting ourselves to species present on that tree (2685 out of 6882 species).

### Linkage between CRISPR array and *cas* genes

Often CRISPR arrays and *cas* genes are collocated such that loss of one may be linked to loss of the other. In a theoretical sense this should not matter, as it will not alter the asymptotic distribution of array counts per genome that we would expect in the case of either functional or non-functional genomes. That is, while linked-loss could lower the number of arrays seen in non-functional genomes in the short term, it should not change the value of the eventual equilibrium array count a genome tends toward over time.

Nevertheless, genomes may not be at or near their equilibrium array counts. We can test this assumption directly by regressing the species-specific values of Δ*µ*_*k*_ (defined above when correcting for lineage-specific trends in HGT) against the minimum distance between CRISPR arrays and *cas* genes on a genome. If CRISPR-*cas* linkage were driving our results, we would see a strong relationship between these values. We include only completely assembled genomes in this analysis as genomic distances are needed.

### CRISPR spacer turnover model

We develop a simple deterministic model of the spacer turnover dynamics in a single CRISPR array of a bacterium exposed to *n* viral species (i.e., disjoint protospacer sets):

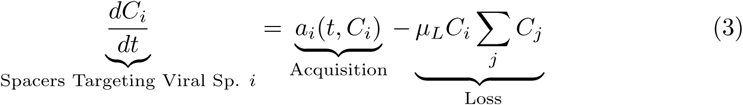

where *µ*_*L*_ is the spacer loss rate parameter and *a*_*i*_ is a function of time representing the viral environment. Here we let *a*_*i*_(*t, C*_*i*_) = *µ*_*A*_*v*_*i*_*f*_*i*_(*t*), where *µ*_*A*_ is the spacer acquisition rate, *v*_*i*_ is a composite parameter describing the maximum density of a viral species in the environment multiplied by the adsorption rate, and *f*_*i*_(*t*) is a function describing the fluctuations of the viral population over time that takes values between zero and one.

The rate of per-spacer loss increases linearly with locus length. This assumption is based on the observation that spacer loss appears to occur via homologous recombination between repeats (Garrett et al., 2011; Gudbergsdottir et al., 2011; Weinberger et al., 2012a), which becomes more likely with increasing numbers of spacers (and thus repeats). Using this model we can determine optimal spacer acquisition rates given a particular pathogenic environment. If there are multiple optima, or if optima cluster in different regions of parameter space for different pathogenic environments, this indicates that having multiple-arrays may be the best solution in a given environment or set of environments.

We analyze a simple case with two viral species where there is one “background” species (*B*) representing the set of all viruses persisting over time in the environment (*f*_*B*_(*t*) = 1) and another “transient” species (*T*) that leaves and returns to the environment after some interval of time (*f*_*T*_ (*t*) is a binary function that takes a value of one if virus *T* is present in the environment and zero otherwise). This allows us to effectively illustrate any tradeoff between the ability to maintain defenses towards a constant threat and the ability to defend against threats that periodically reappear in the environment. In practice we will focus on one interval in which *T* leaves and then returns to the system in order to see if immune memory is lost, and if so how long it takes to regain that memory, given a particular spacer acquisition rate (*µ*_*A*_). We also find how long it would take to acquire immunity towards a completely novel phage species given a particular acquisition rate in order to assess any tradeoff between learning-speed and memory-span.

We can also include the phenomenon of priming in our model, wherein if a CRISPR array has a spacer targeting a particular viral species, the rate of spacer acquisition towards that species is increased (Datsenko et al., 2012; Swarts et al., 2012). Thus

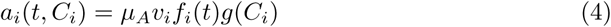

where

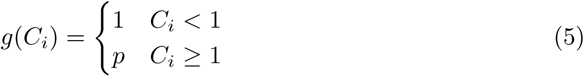

is a stepwise function determining the presence or absence of at least one spacer towards a given viral species and *p* > 1 is the degree of priming. For details of model analysis see S1 Text.

There is evidence that mature crRNA transcripts from the leading end of the CRISPR array are far more abundant than those from the trailing end, and that this decay over the array happens quickly (most transcripts are from the first few spacers, though some spacers farther down can occasionally show high representation) (Bernick et al., 2012; Hale et al., 2012; Richter et al., 2012). This suggests an alternative model wherein the length of an “effective” array is capped at a constant number of spacers and “loss” (i.e. movement out of the zone where crRNAs are actively transcribed and processed) occurs directly due to the acquisition of novel spacers. Therefore we built a modified version of the model above with a hard cap on array length (S1 Text).

## Results

### Having more than one CRISPR array is common

About half of the prokaryotic genomes in the RefSeq database have at least one CRISPR array (44%). Of these genomes, more than half have more than one CRISPR array (63%). When restricting ourselves only to putatively functional genomes where the CRISPR spacer acquisition machinery was present (*cas1* and *cas2*) the proportion of genomes with more than one array increases to 68%. In contrast to this result, having more than one set of *cas* targeting genes is not nearly as common. Signature targeting genes are diagnostic of CRISPR system type. We counted the number of signature targeting genes for type I, II, and III systems in each genome (*cas3, cas9*, and *cas10* respectively Makarova et al. 2015). Only 5% of all genomes have more than one targeting gene (either multiple copies of a single type or multiple types). Even when restricting ourselves again to genomes with a CRISPR array, only 10% of genomes had multiple signature targeting genes. However, of those genomes with more than one set of targeting genes, many had multiple types (48%).

Some taxa are overrepresented in RefSeq (e.g. because of medical relevance), and we wanted to avoid results being driven by just those few particular taxa. We controlled for this by randomly sub-sampling 10 genomes from each taxon with more than 10 genomes in the database and found broadly similar results. After sub-sampling, approximately 40% of genomes had at least one CRISPR array, and of these 61% had more than one. Of genomes with intact spacer acquisition machinery, 62% had more than one CRISPR array. Of these sub-sampled genomes, restricting to those with at least one CRISPR array, 9% had more than one set of *cas* targeting genes. Of these multi-*cas* genomes, many had multiple types (56%).

### Validation of functional / non-functional classification

Our power to detect selection depends critically on our ability to classify genomes as CRISPR functional vs. non-functional. Functional CRISPR arrays should, on average, contain more spacers than non-functional arrays (Gophna et al., 2015). Thus we compared the number of repeats in CRISPR arrays in genomes with both *cas1* and *cas2* present (“functional”, 16.01 repeats on average) to the number of spacers in genomes lacking either or both genes (“non-functional”, 12.23 repeats on average) and confirmed that the former has significantly more than the latter (*t* = −36.516, *df* = 55340, *p <* 2.2 × 10^−16^; S1 Fig). This difference in length (3.80 repeats) is not as large as one might expect, possibly because some systems are able to acquire or duplicate spacers via homologous recombination (Kupczok et al., 2015) and arrays may have been inherited recently from strains with active *cas* machinery. The mean array length across the dataset was 15.12 repeats.

### Instantaneous array loss vs. gradual decay

There are two possible routes to complete CRISPR array loss: (1) an all-at-once loss of the array (e.g. due to recombination between flanking insertion sequences Almendros et al. 2014; Shah and Garrett 2011) and (2) gradual decay due to spacer loss. Limited experimental evidence supports (1) spontaneous loss of the entire CRISPR array (Jiang et al., 2013), as do comparisons between closely related genomes (Shah and Garrett, 2011). The distinction above is important, because if CRISPR array loss were to occur primarily via (2) gradual decay, then functional genomes would have an intrinsically lower rate of array loss than non-functional genomes. This is because in functional genomes spacer acquisition would counteract spacer loss, reducing the rate of array decay, whereas this compensation would not occur in non-functional genomes. This could lead us to spuriously accept a result of selection maintaining multiple arrays.

If arrays were primarily lost via gradual decay we would expect a positive relationship between the number of arrays in a genome and the average array length in a genome, because arrays experiencing more decay (either due to increased spacer loss rates or reduced acquisition rates) should be shorter and prone to eventual deletion. In functional genomes with the complete spacer acquisition machinery (*cas1* and *cas2*) this trend would be due to the higher probability of stochastically reaching a 0-spacers state in shorter arrays, and arrays will in general be shorter in genomes with lower spacer acquisition rates. In non-functional genomes that lack the complete spacer acquisition machinery, this trend would result from differences in time since loss of acquisition machinery, where genomes that had lost that machinery farther in the past would have both shorter arrays and fewer arrays on average.

Overall we see no relationship between mean array length and array count in a genome (*m* = −0.001, *p* = 0.109, *R*^2^ = 5.55×10^−5^). Surprisingly, in functional genomes we find a slightly negative linear relationship between mean array length in a genome and the number of arrays in a genome (*m* = −0.0081, *p <* 2×10^−16^, *R*^2^ = 0.0032). In non-functional genomes we see a slightly positive relationship (*m* = 0.0054, *p* = 7.23×10^−10^, *R*^2^ = 0.0026). While both of these relationships are significant, they are extremely weak and probably spurious. Thus we reject any clear array-length vs. array count relationship and accordingly rule out array loss via spacer-wise decay as a major driver of the patterns we will explore later.

### Selection maintains multiple CRISPR arrays

We leveraged the difference between functional and non-functional genomes, within each of which the process of CRISPR array accumulation should be distinct (Fig. 1, Table 1). Non-functional CRISPR arrays should accumulate neutrally in a genome following background rates of horizontal gene transfer and gene loss. We constructed two point estimates of this background accumulation process using our parametric model to infer the distribution of the number of arrays. One estimate came directly from the non-functional genomes (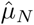, Fig. 1a). The other came from the functional genomes, assuming that having one array is adaptive in these genomes, but that additional arrays accumulate neutrally (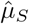, Fig. 1b). If selection maintains multiple (functional) arrays, then we should find that 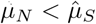. We found this to be overwhelmingly true, with about two arrays on average seeming to be evolutionarily maintained across prokaryotic taxa 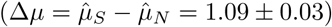. We bootstrapped 95% confidence intervals of our estimates (Table 1) and found that the bootstrapped distributions did not overlap, indicating a highly significant result (Fig. 1d)

**Table 1.** Tests for multi-array adaptiveness applied to different subsets of the RefSeq data. See Fig 1 and S2 Fig-S4 Fig.

| Only $\leq 1$ *cas* set | Sub-sampled | $\hat{\mu}_S$ | Bootstrap | | $\hat{\mu}_N$ | Bootstrap | | $\Delta\mu$ | $s^*$ |
| --- | --- | --- | --- | --- | --- | --- | --- | --- | --- |
|  |  |  | 2.5% | 97.5% |  | 2.5% | 97.5% |  |  |
| No | No | 1.56 | 1.53 | 1.58 | 0.46 | 0.45 | 0.47 | 1.09 | 2 |
| No | Yes | 2.26 | 2.19 | 2.34 | 1.13 | 1.08 | 1.17 | 1.14 | 2 |
| Yes | No | 1.05 | 1.04 | 1.07 | 0.45 | 0.44 | 0.46 | 0.61 | 2 |
| Yes | Yes | 1.26 | 1.19 | 1.32 | 1.07 | 1.03 | 1.12 | 0.19 | 1 |

**Fig 1.**
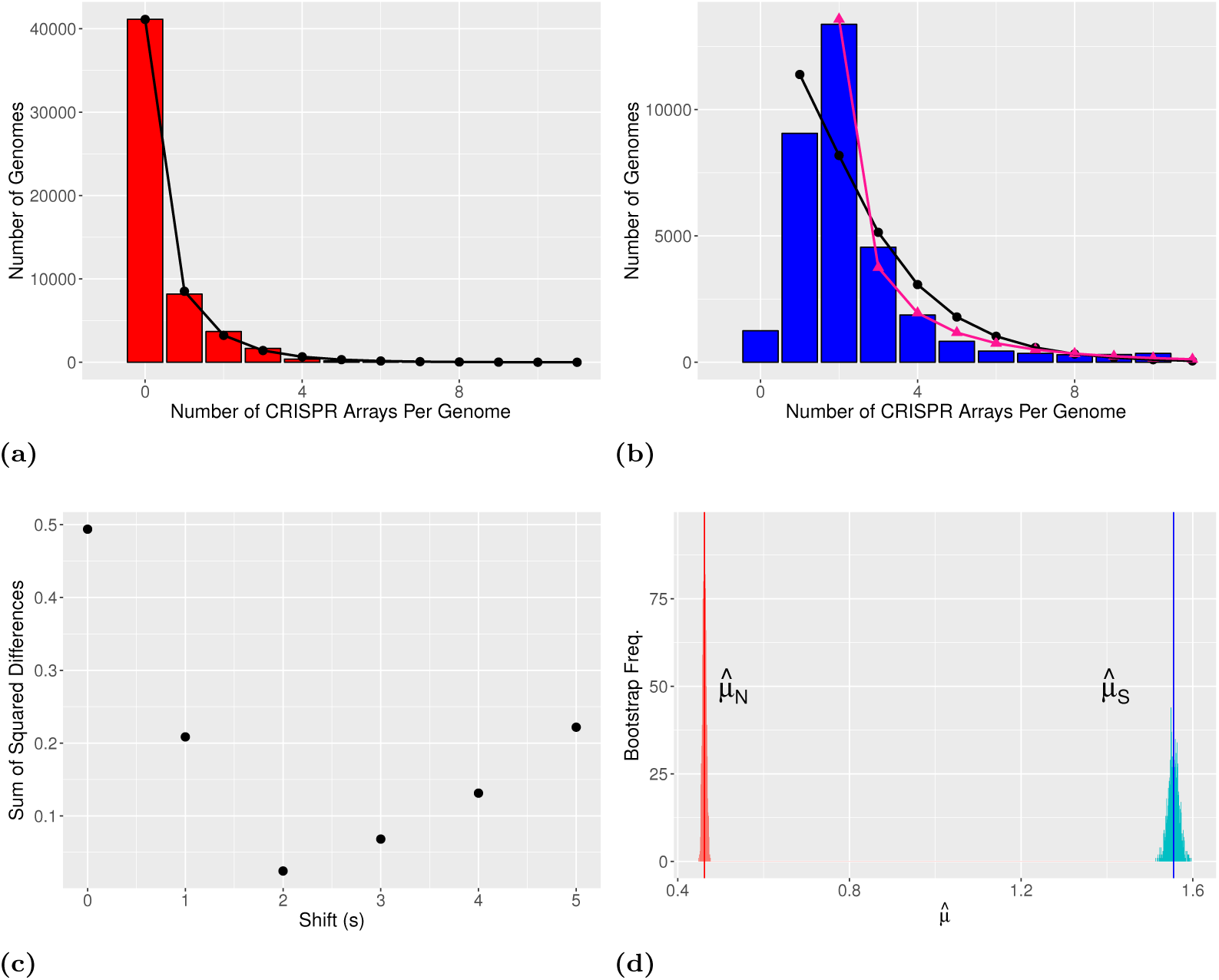
Selection maintains more than one CRISPR array on average across prokaryotes. (a-b) Distribution of number of arrays per genome in (a) genomes with non-functional CRISPR immunity and (b) genomes with putatively functional CRISPR immunity. The tails of these distributions are cut off for ease of visual comparison (24 genomes with > 10 arrays in (a) and 498 genomes with > 10 arrays in (b)). In (a) the black circles show the negative binomial fit to the distribution of arrays in non-functional genomes. In (b) black circles indicate the negative binomial fit to the single-shifted distribution (*s* = 1) and pink triangles to the double-shifted distribution (*s* = 2). (c) The optimal shift is *s*\* = 2, where the difference between the two distributions is minimized. (d) The bootstrapped distributions of the parameter estimates of 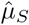 and 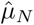 show no overlap with 1000 bootstrap replicates.

Sub-sampling overrepresented taxa altered our parameter estimates slightly, but did not change our overall result (Δ*µ* = 1.13±0.09, S2 Fig). To control for the possibility that multiple sets of *cas* genes in a small subset of genomes could be driving this selective signature, we restricted our dataset only to genomes with one or fewer signature targeting genes (*cas3, cas9*, or *cas10* Makarova et al. 2011a, 2015) *and one or fewer copies each of the genes necessary for spacer acquisition (cas1* and *cas2*). Even when restricting our analyses to genomes with one or fewer sets of *cas* genes, there is selection to maintain more than one (functional) CRISPR array, though the effect size is smaller (Δ*µ* = 0.61 *±* 0.02, S3 Fig; with sub-sampling of overrepresented taxa Δ*µ* = 0.19 *±* 0.08, S4 Fig).

### Confirming selection

The number of CRISPR arrays is positively related to the number of *cas* genes in a genome (S5 Fig). To control for the potentially confounding effect of variation in the rate of HGT among lineages, we performed several additional analyses. First, if we further restrict our sub-sampled dataset of genomes with one or fewer sets of *cas* genes, such that each species is represented by an equal number of functional and non-functional genomes, then we still find a signature of selection on multiple arrays (Δ*µ* = 0.40±0.15, S6 Fig). Unfortunately this method involves excluding a large portion of the dataset. Second, our species-wise implementation of the Δ*µ* test that controls for differences in rates of HGT between lineages also confirms a signature of multi-array selection 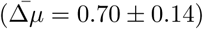. Because there is a low number of genomes for most species and this test restricts us to only within-species comparisons, our species-wise parameter-based suffers somewhat in terms of power.

Third, we fit a linear model accounting for phylogeny (Tung Ho and Anè (2014)) of the mean number of CRISPR arrays on a genome per species versus the proportion of functional genomes per species. This yielded a slope near two, indicating maintenance of two arrays on average (*m* = 1.9283, *p <* 2×10^−16^), and was actually higher than the slope found not considering phylogeny (*m* = 1.69788, *p <* 2×10^−16^).

In order to determine if the signature of selection on multiple arrays we observed was confined to a particular set of clades, we mapped all species-specific Δ*µ*_*k*_ values onto the SILVA Living Tree 16s rRNA tree (Yarza et al., 2008). Of the 623 species with at least one functional and one non-functional genome, 568 were represented on the tree. Positive and strongly positive (> 1) values of Δ*µ*_*k*_ were distributed across the tree, indicating this phenomenon was not isolated to a particular group (S12 Fig). Formal testing revealed no significant phylogenetic signal in the Δ*µ*_*k*_ values (*K* = 1.88×10^−9^, *p* = 0.604; Blomberg et al. 2003; Revell 2012)

We also regressed the species-specific Δ*µ*_*k*_’s found above against the minimum distance between CRISPR arrays and *cas* genes in a genome to verify that linkage was not driving our result. We saw a slight positive relationship between CRISPR-*cas* distance and our signature of multi-array selection, the opposite of what we would expect if linkage were driving our results (*m* = 3.163×10^−7^, *p* = 8.52×10^−6^, *R*^2^ = 0.009937).

Finally, to confirm that assembly level had no effect on our outcome, we ran our parametric test restricted to completely assembled genomes in the dataset. This reduced dataset still yielded a signature of selection on multiple array (6263 genomes, Δ*µ* = 0.98 *±* 0.09)

### Neo-CRISPR Arrays

Recently, Nivala et al. (2018) found that off-target spacer integration into the genome can spawn novel CRISPR arrays in *E. coli*. This could create a spurious signature of selection maintaining multiple arrays using our test, since the production of “neo-CRISPR arrays” would only occur in functional genomes. A simple way to control for this is to merge all CRISPR arrays with identical consensus repeat sequences in a genome, thus removing any duplicates. Doing this, we find that the signature of multi-array selection remains, albeit being somewhat less strong (Δ*µ* = 0.46±0.02). We were considerably surprised that this signature of selection still remained after merging, since such merging will also remove a large portion of arrays acquired through horizontal transfer, assuming such transfers most often happen between closely related individuals. In any case, while the production of neo-CRISPR arrays may be driving our result in part, it cannot account for the overall signal. It is unclear if neo-CRISPR arrays are commonly produced in bacteria via off-target integration, though Nivala et al. (2018) found circumstantial evidence it may occur in two other species. The CRISPR system of *E. coli* is not naturally active (Savitskaya et al., 2017) and requires artificial up-regulation of the spacer acquisition machinery.

### Validation of CRISPRDetect array predictions

We ran our tests for selective maintenance of multiple arrays on the same dataset excluding arrays with a CRISPRDetect score lower than 6 (double the default threshold). We found no qualitative differences in our results when we used this increased detection threshold (Δ*µ* = 1.00±0.02). By default, CRISPRDetect identifies arrays with repeats matching experimentally-verified CRISPR arrays as well as *de novo* repeats. If we restrict to only arrays with a positive hit on this list we again found the same pattern (Δ*µ* = 0.76±0.03).

We also downloaded the set of CRISPR arrays and array-lacking genomes available on the CRISPR Database (Grissa et al., 2007a). This database uses an alternative algorithm for array detection (Grissa et al., 2007b) and thus serves as an independent verification of our results. This dataset showed a clear signature of selection maintaining multiple arrays (Δ*µ* = 1.49 *±* 0.17).

### Evidence for array specialization

In genomes with multiple arrays, the dissimilarity between consensus repeat sequences of arrays in a single genome spanned a wide range of values (Levenshtein Distance, S7 Fig and S8 Fig), though the mode was at zero (i.e., identical consensus repeats). When limiting our scope to only genomes with exactly two CRISPR arrays, we saw a bimodal distribution of consensus repeat dissimilarity, with one peak corresponding to identical arrays within a genome and the other corresponding to arrays with essentially randomly drawn repeat sequences except for a few conserved sites between them (S7D Fig). We also observed that among functional genomes, the area of the peak corresponding to dissimilar repeat sequences was significantly higher than among non-functional genomes (*χ*^2^ = 61.432, *df* = 1, *p <* 4.582 *×* 10^−15^, S7 Fig). This suggests that the observed signature of adaptiveness may be related to the diversity of consensus repeat sequences among CRISPR arrays in a genome.

We attempted to independently confirm this result using repeat sequences obtained from the CRISPR Database. While the distribution of pairwise-distances between repeat sequences in two-array genomes was approximately the same shape as that we observed for our dataset (S8 Fig), the relationship between diversity and functionality was reversed, with non-functional genomes having more diverse consensus repeats among their arrays (*χ*^2^ = 4.3952, df = 1, *p* = 0.03604). This opposing result calls into question the pattern observed in the CRISPRDetect data, though this may be due to the smaller size of the CRISPR Database dataset.

### A tradeoff between memory span and acquisition rate could select for multiple arrays in a genome

We hypothesized that having multiple systems with different acquisition rates could allow prokaryotes to respond to a range of pathogens with different characteristics (e.g. residence time in the environment, frequency of recurrence). To investigate this possibility we built a simple model of spacer turnover dynamics in a single CRISPR array in the presence of “background” and “transient” viral species (see Methods). We constructed phase diagrams of the model behavior, varying spacer acquisition rates and the relative population sizes of viral species or the extent of priming, respectively (Fig. 2a, S9 Fig). We found that for very high spacer acquisition rates, the system is able to maintain immunity to both background and transient viral populations (“short-term memory”/”fast-learning”). High rates of spacer acquisition are unrealistic as they lead to high rates of autoimmunity (S2 Text, Wei et al. 2015; Kumar et al. 2015; Yosef et al. 2012; Levy et al. 2015; Stern et al. 2010). Our analysis also reveals that there is a region of parameter space with low spacer acquisition rates in which immunity is maintained (“long-term memory”/”slow-learning”). This is the region where low spacer turnover rates allow immune memory to remain in the system over longer periods of time (Fig. 2a).

**Fig 2.**
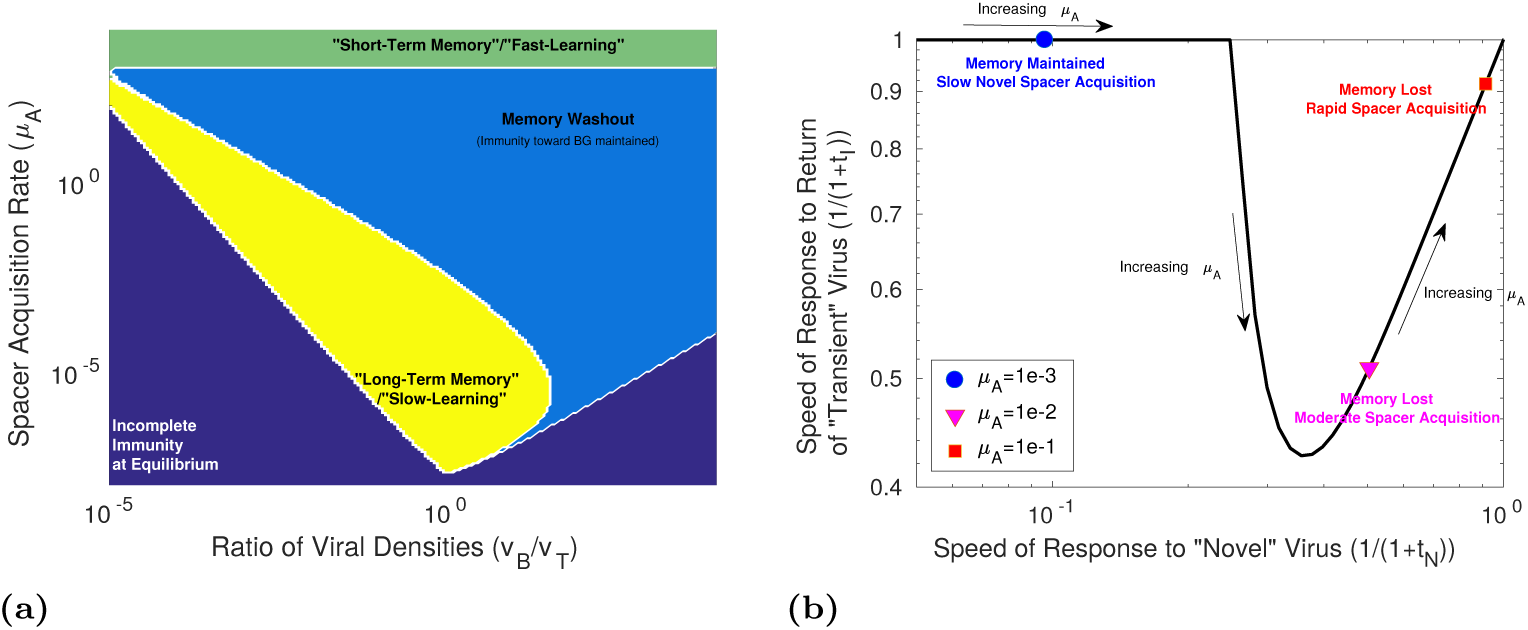
Immune memory is maximized at intermediate and low spacer acquisition rates, creating a tradeoff with the speed of immune response to novel threats. (a) Phase diagram of the behavior of our CRISPR array model with two viral species, a constant “background” population and a “transient” population that leaves and returns to the system at some fixed interval. The yellow region indicates that immunity towards both viral species was maintained. The green region indicates where immune memory was lost towards the transient phage species, but reacquired almost immediately upon phage reintroduction (*t*_*I*_ *<* 10^−5^, where *t*_*I*_ is the time to first spacer acquisition after the return of the species to the system following an interval of absence). The light blue region indicates that only immunity towards the background species was maintained (i.e., immune memory was rapidly lost and *t*_*I*_ > 10^−5^). Dark blue indicates where equilibrium spacer content towards one or both species did not exceed one despite both species being present in the system (S1 Text). (b) The tradeoff between memory span and learning speed. The speed of immune response to the transient viral species (as in (a), with *v*_*B*_*/v*_*F*_ = 1) is plotted against the speed of response to a novel viral species to which the system has not been previously exposed (so that there are no spacers targeting this species), over a range of *µ*_*A*_ values (*µ*_*A*_ *∈* [10^−3.5^, 1]). The speed of immune response to the transient species is defined as 1*/*(1 + *t*_*I*_) (where *t*_*I*_ = 0 if memory is maintained). The speed of response to the novel species is similarly defined as 1*/*(1 + *t*_*N*_) where *t*_*N*_ is the time to first spacer acquisition towards this species. For specifics on calculating *t*_*I*_ and *t*_*N*_ see S1 Text.

The “long-term memory”/”slow-learning” region of parameter space is separated from the “short-term memory”/”fast-learning” region of parameter space by a “memory-washout” region in which spacer turnover is high so that memory is lost but acquisition is not rapid enough to quickly re-acquire immunity towards the transient virus (Fig. 2a). The relative densities of the different viral species modulate the relative importance of fast-acquisition versus memory span. Thus for a range of pathogenic environments two distinct CRISPR immune strategies exist with respect to the spacer acquisition rate (“long-term memory” vs. “fast-learning”). We also note that high levels of priming expand the “washout” region separating the two strategies, as high spacer uptake from background viruses will crowd out long term immune memory (S9 Fig).

A single CRISPR array fulfilling one of the two CRISPR strategies is sufficient in the case shown in Fig 2a, although only one of those strategies is likely accessible due to limits on spacer acquisition rates. When a third, novel viral species enters the system, a two-array solution may become necessary due to the memory span vs. learning speed tradeoff (Fig 2b). Extremely high spacer acquisition rates might allow the host to rapidly respond to both novel and returning threats, but, as noted above, such rates are unrealistic due to physical constraints on the speed of adaptation as well as the evolutionary constraint of autoimmunity (S2 Text, Wei et al. 2015; Kumar et al. 2015; Yosef et al. 2012; Levy et al. 2015; Stern et al. 2010). CRISPR adaptation is rapid, but it is not instantaneous, and infected hosts with arrays lacking an appropriate spacer will often perish before a spacer can be acquired (Hynes et al., 2014). This means that the region of rapid immune response to both transient and novel threats by a single array, shown in the top-right corner of Fig 2b, is probably inaccessible. Thus, in order to maximize novel spacer acquisition and memory span simultaneously, a two-system solution will be required.

An alternative model with fixed-length arrays, corresponding to a situation in which only the first few spacers in an array are processed into mature crRNAs (Bernick et al., 2012; Hale et al., 2012; Richter et al., 2012), demonstrated qualitatively similar behavior to the model described above (S1 Text, S10 Fig).

### Taxon-specific signatures of selection

Several taxa in the dataset were represented by a large number of genomes (> 1000), with at least one each of functional and non-functional genomes. We performed our test for adaptiveness on each of these taxa individually (Table 2). We found that among *Staphylococcus aureus, Klebsiella pneumoniae, Shigella sonnei*, and *Listeria monocytogenes* genomes there was a strong signal of selection maintaining multiple arrays.

**Table 2.** Species specific values of Δ*µ*

| Species | $\Delta\mu$ |
| --- | --- |
| *Staphylococcus aureus* | $1.15 \pm 0.37$ |
| *Klebsiella pneumoniae* | $0.76 \pm 0.06$ |
| *Shigella sonnei* | $0.72 \pm 0.17$ |
| *Listeria monocytogenes* | $0.67 \pm 0.08$ |
| *Mycobacterium tuberculosis* | $0.41 \pm 0.05$ |
| *Pseudomonas aeruginosa* | $0.35 \pm 0.16$ |
| *Campylobacter jejuni* | $-0.12 \pm 0.05$ |
| *Escherichia coli* | $-0.20 \pm 0.04$ |
| *Salmonella enterica* | $-0.54 \pm 0.06$ |

Surpisingly, genomes of *Campylobacter jejuni, Eschericia coli*, and *Salmonella enterica* show evidence for selection against having a CRISPR array in the presence of functional *cas* machinery. Previous work has shown that CRISPR in *E. coli* and *S. enterica* appears to be non-functional as an immune system under natural conditions (Touchon and Rocha, 2010; Touchon et al., 2011). All of these taxa are human pathogens, and can occupy a diverse set of environmental niches on the human body. It is unclear at this time what is causing the differences in the adaptive landscape each taxon experiences.

A very small portion of the genomes used in our analyses were from archaea (*<* 1%). We ran our analyses on these genomes alone to see if they differed qualitatively from their bacterial counterparts. Archaeal genomes showed a clear signal of selection maintaining multiple arrays, although the confidence interval around our statistic is rather broad (Δ*µ* = 1.05±0.56). We note that the large majority of archaeal genomes had CRISPR arrays and were also functional, making our approach less powerful (S11 Fig). Further, if those few non-functional genomes lost their *cas* spacer acquisition machinery recently, then our power would be reduced even more because these genomes might still bear the remnants of past selection.

### Type-specific signatures of selection

While we do not have direct information on system type for the majority of arrays in our dataset, we can subdivide genomes into those containing the signature cas targeting genes for type I,II, or III CRISPR systems (*cas3, cas9*, and *cas10* respectively), assuming that this is a reliable indicator of system type (Makarova et al., 2015). The number of arrays per genome differed significantly between each of these pairs (S13 Fig), though the largest difference was between genomes with class I targeting proteins which had around 2 arrays on average (type I and type III, 2.10 and 1.96 respectively) and class II targeting proteins which only had one array on average (type II, 1.05). We excluded genomes with multiple types of targeting genes for this analysis.

We cannot run our test for selection directly on these subsets of the data, since they exclude genomes without arrays or *cas* genes. Instead we classified species into types if the only observed targeting gene type among all representatives of that species corresponded to a a particular type. Thus we can test for our signature of selection among species that “favor” a particular type of CRISPR system. All types showed a signature of multi-array selection (Δ*µ* = 1.09±0.05, 0.62±0.02, 1.79±0.06 respectively). In particular type III “species” had a particularly strong signal, and organisms in this group may be under selection to maintain three arrays.

## Discussion

### Selection maintains multiple CRISPR arrays across prokaryotic taxa

On average, prokaryotes are under selection to maintain more than one CRISPR array. This surprising result holds controlling for both overrepresented taxa and the influence of multiple sets of *cas* genes. However, the degree of selection appears to vary between taxa, likely as a function of the pathogenic environment each experiences based on its ecological niche.

The number of CRISPR arrays in a genome appears to follow a negative binomial distribution quite well (Figs 1b and 1a, S2 Fig-S4 Fig), consistent with our theoretical prediction. This pattern is robust to sub-setting of the data in a variety of ways. We note that, due to the large size of this dataset, formal goodness-of-fit tests to the negative binomial distribution always reject the fit due to small but statistically significant divergences from the theoretical expectation.

Our test for selection is conservative to the miscategorization of arrays as “functional”or “non-functional”. Miscategorizations could occur because intact targeting machinery still allows for preexisting spacers to confer immunity, some CRISPR arrays may be conserved for non-immune purposes (e.g. Touchon and Rocha 2010; Li et al. 2016), or intact acquisition machinery is no guarantee of system functionality. That being said, our test is conservative precisely because of such miscategorizations, as they should increase 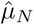 and decrease 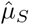 respectively. Selection against having a CRISPR array in genomes lacking spacer acquisition machinery could produce a false signature of selection maintaining arrays in genomes with acquisition machinery. This is unlikely because there is no reason a non-functional CRISPR array should be under strong negative selection given the low or nonexistent associated costs.

Our test should also be robust to false positive or negative array discovery rates. Because we used CRISPRDetect settings such that *cas*-gene presence was not taken into account when scoring arrays, CRISPRDetect followed identical procedures for detecting arrays in functional and non-functional genomes. Thus an elevated false-positive or false-negative rate should have no effect on our tests for selection, because these tests rely on relative differences between array counts in functional and non-functional genomes, not their actual values. We further confirmed this by changing our CRISPRDetect score threshold and comparing to the distribution of arrays per genome in the CRISPR Database (Grissa et al., 2007a).

Finally, we note that *µ*_*N*_ and *µ*_*S*_ take on a range of values depending on what subset of taxa/genomes is considered. This is to be expected as each set of species will occupy a distinct environment in terms of both the rate of horizontal gene transfer and the usefulness of CRISPR immunity. Nevertheless, our qualitative result of multi-CRISPR adaptiveness is robust to this quantitative variability.

### Why have multiple CRISPR-Cas systems?

Possibly, multiple arrays could be selectively maintained even in the absence of any fitness advantage if each array acquired complementary spacer content towards distinct viral targets by chance. If arrays share acquisition machinery such complementarity is unlikely because priming will ensure both arrays contain spacers towards any target encountered, meaning that the content of the two arrays will be largely redundant. We note that this argument only holds for type I and II systems which demonstrate priming (Datsenko et al., 2012). Type III systems are unprimed, have slow spacer acquisition rates, and can target even mutated viral sequences (Pyenson et al., 2017). Thus type III systems may be maintained via spacer complementarity, perhaps explaining why species favoring type III systems appear to experience selection maintaining three rather than just two CRISPR arrays.

Even in type I and II systems, if each array is associated with a separate set of spacer acquisition machinery, then cross-priming will be less likely and complementarity could arise. Nevertheless, this does not explain the multi-array conservation we see in genomes with only a single set of cas genes. Thus we consider here several potential adaptive explanations for the selection that maintains multiple arrays.

Our data show significant numbers of both functionally similar and dissimilar CRISPR systems within the same genome, so either could potentially be adaptive. While CRISPR systems are generally highly flexible, a prokaryote might still gain an advantage in the former case if multiple similar systems lead to improved immunity through redundancy and in the latter case if multiple dissimilar systems allow for specialization towards multiple types of threats. The relevance of the different advantages depends on whether an individual has multiple sets of *cas* genes, CRISPR arrays, or both.

In the case of similar systems, immunity could be improved by (a) an increased spacer acquisition rate, (b) an increased rate of targeting, or (c) a longer time to expected loss of immunity. Duplication of *cas* genes could, in principle, increase uptake (a) and targeting rates (b) through increased gene expression, but our data show that multiple sets of *cas* genes are rare, which suggests this is, at best, a minor force. Alternatively, duplication of CRISPR arrays could increase targeting (b) via an increased number of crRNA transcripts or increase memory duration (c) through spacer redundancy. However, the effectiveness of crRNA may actually decrease in the presence of competing crRNAs (Stachler and Marchfelder, 2016; Stachler et al., 2017) and, since a single array can have multiple spacers with the same target, there is a diminished advantage to having multiple arrays in terms of memory span (S3 Text). Redundant arrays might also be a form of bet-hedging since CRISPR functionality is lost at a high rate in some prokaryotes (Jiang et al., 2013; Weissman et al., 2018).

In the case of dissimilar systems, immunity could be aided if diverse features are advantageous. For example some viruses encode proteins that deactivate Cas targeting proteins (Bondy-Denomy et al., 2013; Pawluk et al., 2014; Rauch et al., 2017). Diverse *cas* genes may allow hosts to evade the action of these anti-CRISPR proteins, which are often extremely broadly acting (Bondy-Denomy et al., 2013; Pawluk et al., 2014). Alternatively, it has recently been shown that promiscuous type III Cas proteins are often encoded alongside type I systems and can function as a “backup”, using spacers from the same array to target phages that have mutated the protospacer-adjacent motifs necessary for type I targeting (Silas et al., 2017). Many genomes with multiple *cas* signature genes also had multiple types of such genes, possibly indicating some diversifying force. The inclusion of these multi-*cas* genomes also increased the effect size of our test for adaptiveness, despite low representation in the dataset. In any case, while these hypotheses remain interesting candidates to explain CRISPR multiplicity in some prokaryotes, the majority of the genomes in the dataset have only one set of *cas* genes and thus these mechanisms cannot explain the signature for multi-array adaptiveness observed in the majority of the dataset.

The theoretical model we present demonstrates one possible explanation for selective maintenance of multiple CRISPR arrays. Rate variation is an attractive hypothesis since it can explain the signature of selective maintenance we observe even in the absence of multiple sets of cas genes. Additionally, rates vary between systems, and rate variation among arrays on a single genome has been observed in a diverse set of organisms (e.g. (Paez-Espino et al., 2015; Westra et al., 2015)). Arrays with slightly different, or even identical, consensus repeat sequences may differ in length, despite sharing a set of *cas* genes (Zeng et al., 2017). It has even been shown directly that acquisition rates can vary among CRISPR arrays with identical consensus repeat sequences sharing a single set of *cas* genes (Staals et al., 2016). The factors influencing acquisition rate appear to be idiosyncratic, perhaps related to the genomic position of the CRISPR array.

We do not provide empirical evidence that rate variation drives the observed signature of selection of multiple arrays. Such verification is not currently possible, and will require the characterization of spacer acquisition and loss rates across a large number of taxa. Nevertheless, we develop this hypothesis as a promising candidate and illustrate how factors intrinsic to the mechanism of CRISPR immunity can create strong tradeoffs between memory span and learning speed. We show how such tradeoffs can lead to selection for both high acquisition rate (i.e., short term memory) and low acquisition rate (i.e., long-term memory) systems, depending on the pathogenic environment of the host. As an array increases in length (i.e., the number of repeats increases) the rate of spacer loss should also increase because loss occurs via homologous recombination. Thus a length-dependent spacer loss rate causes high acquisition rate systems to also have high loss rates, producing the aforementioned tradeoff.

Our primary model demonstrates that array-length dependent spacer loss can produce a “learning” versus “memory” tradeoff. Nevertheless, in some systems spacer loss may occur at a very low rate, making our proposed mechanism less relevant. There is some evidence of spacers that persist in a population over a timespan of years (Tyson and Banfield, 2008), though such conservation is possibly due to selection on the population rather than a low mechanistic loss rate. A slightly modified model that imposes a hard cap on the number of “effective” spacers in an array (S1 Text) and would not require a strong spacer loss vs. acquisition rate link produced qualitatively similar results. We based this model on evidence that the abundance of mature crRNA transcripts from the leader end greatly exceeds that of those from the trailing end (Bernick et al., 2012; Hale et al., 2012; Richter et al., 2012), implying that only the first few spacers in an array will provide effective immunity. Thus CRISPR immunity may face a memory-learning tradeoff due to multiple, distinct mechanisms, but in both cases spacer acquisition rate variation among systems can lead to an optimal solution for the host.

We note that when partial spacer-target matches exist, variability in spacer acquisition rates among arrays will be largely irrelevant since priming will ensure rapid acquisition of new spacers. On the other hand, when no match exists, either due to spacer loss or the introduction of a truly novel viral species into the environment, primed spacer uptake will not occur. Thus the rate at which a host encounters novel threats will determine the relative importance of the baseline spacer acquisition rate versus the primed acquisition rate. In environments where novel viruses are frequently encountered, small differences in acquisition rate can be important for host fitness, whereas in environments where host and virus pairs consistently coevolve over time priming will be the more important phenomenon.

As more genome sequences from environmental samples become available, it will be possible to explicitly link particular array configurations to specific features of the pathogenic environment or host lifestyle. Even then, open questions remain. One phenomenon that we do not address here is that a small, but non-trivial number of genomes have greater than 10 arrays. It is difficult to imagine so many arrays accumulating neutrally in a genome. If high array counts are a product of high horizontal transfer rates, then genomes with extremely high array counts should also be larger due to accumulation of foreign genetic material. This was not the case (S14 Fig), indicating that rates of horizontal transfer alone cannot explain these outliers.

Finally, our examination of immune configuration is likely relevant to the full range of prokaryotic defense mechanisms. In contrast to previous work focusing on mechanistic diversity (e.g. Iranzo et al. 2013, 2015; Kumar et al. 2015; Westra et al. 2015), we emphasize the importance of the multiplicity of immune systems in the evolution of host defense. As we suggest, a surprising amount of strategic diversity may masquerade as simple redundancy.

## Acknowledgments

JLW was supported by a GAANN Fellowship from the U.S. Department of Education and the University of Maryland. WFF was partially supported the U.S. Army Research Laboratory and the U.S. Army Research Office under Grant W911NF-14-1-0490. PLFJ was supported in part by NIH R00 GM104158.

## References

Almendros C, Mojica FJ, Díez-Villaseñor C, Guzmán NM, García-Martínez J. 2014. Crispr-cas functional module exchange in escherichia coli. MBio. 5:e00767–13.

Andersen JM, Shoup M, Robinson C, Britton R, Olsen KEP, Barrangou R. 2016. CRISPR Diversity and Microevolution in Clostridium difficile. Genome Biology and Evolution. 8:2841–2855.

Barrangou R, Fremaux C, Deveau H, Richards M, Boyaval P, Moineau S, Romero DA, Horvath P. 2007. CRISPR Provides Acquired Resistance Against Viruses in Prokaryotes. Science. 315:1709–1712.

Bernick DL, Cox CL, Dennis PP, Lowe TM. 2012. Comparative genomic and transcriptional analyses of CRISPR systems across the genus Pyrobaculum. Frontiers in Microbiology. 3.

Biswas A, Staals RH, Morales SE, Fineran PC, Brown CM. 2016. CRISPRDetect: A flexible algorithm to define CRISPR arrays. BMC Genomics. 17:356.

Blomberg SP, Garland Jr T, Ives AR. 2003. Testing for phylogenetic signal in comparative data: behavioral traits are more labile. Evolution. 57:717–745.

Bolotin A, Quinquis B, Sorokin A, Ehrlich SD. 2005. Clustered regularly interspaced short palindrome repeats (CRISPRs) have spacers of extrachromosomal origin. Microbiology. 151:2551–2561.

Bondy-Denomy J, Pawluk A, Maxwell KL, Davidson AR. 2013. Bacteriophage genes that inactivate the CRISPR/Cas bacterial immune system. Nature. 493:429–432.

Boudry P, Semenova E, Monot M, et al. (11 co-authors). 2015. Function of the CRISPR-Cas System of the Human Pathogen Clostridium difficile. mBio. 6:e01112–15.

Burstein D, Harrington LB, Strutt SC, Probst AJ, Anantharaman K, Thomas BC, Doudna JA, Banfield JF. 2017. New CRISPR–Cas systems from uncultivated microbes. Nature. 542:237–241.

Burstein D, Sun CL, Brown CT, Sharon I, Anantharaman K, Probst AJ, Thomas BC, Banfield JF. 2016. Major bacterial lineages are essentially devoid of CRISPR-Cas viral defence systems. Nature Communications. 7:10613.

Cai F, Axen SD, Kerfeld CA. 2013. Evidence for the widespread distribution of CRISPR-Cas system in the Phylum Cyanobacteria. RNA Biology. 10:687–693.

Datsenko KA, Pougach K, Tikhonov A, Wanner BL, Severinov K, Semenova E. 2012. Molecular memory of prior infections activates the CRISPR/Cas adaptive bacterial immunity system. Nature Communications. 3:945.

Garrett RA, Shah SA, Vestergaard G, Deng L, Gudbergsdottir S, Kenchappa CS, Erdmann S, She Q. 2011. CRISPR-based immune systems of the Sulfolobales: complexity and diversity. Biochemical Society Transactions. 39:51–57.

Gophna U, Kristensen DM, Wolf YI, Popa O, Drevet C, Koonin EV. 2015. No evidence of inhibition of horizontal gene transfer by CRISPR–Cas on evolutionary timescales. The ISME Journal. 9:2021–2027.

Goren M, Yosef I, Edgar R, Qimron U. 2012. The bacterial CRISPR/Cas system as analog of the mammalian adaptive immune system. RNA biology. 9:549–554.

Grissa I, Vergnaud G, Pourcel C. 2007a. The crisprdb database and tools to display crisprs and to generate dictionaries of spacers and repeats. BMC bioinformatics..

Grissa I, Vergnaud G, Pourcel C. 2007b. CRISPRFinder: a web tool to identify clustered regularly interspaced short palindromic repeats. Nucleic Acids Research. 35:W52–57.

Gudbergsdottir S, Deng L, Chen Z, Jensen JVK, Jensen LR, She Q, Garrett RA. 2011. Dynamic properties of the Sulfolobus CRISPR/Cas and CRISPR/Cmr systems when challenged with vector-borne viral and plasmid genes and protospacers. Molecular Microbiology. 79:35–49.

Hale CR, Majumdar S, Elmore J, et al. (11 co-authors). 2012. Essential features and rational design of CRISPR RNAs that function with the Cas RAMP module complex to cleave RNAs. Molecular Cell. 45:292–302.

Horvath P, Coûté-Monvoisin AC, Romero DA, Boyaval P, Fremaux C, Barrangou R. 2009. Comparative analysis of CRISPR loci in lactic acid bacteria genomes. International Journal of Food Microbiology. 131:62–70.

Houte Sv, Buckling A, Westra ER. 2016. Evolutionary Ecology of Prokaryotic Immune Mechanisms. Microbiology and Molecular Biology Reviews. 80:745–763.

Hynes AP, Villion M, Moineau S. 2014. Adaptation in bacterial CRISPR-Cas immunity can be driven by defective phages. Nature Communications. 5:4399.

Iranzo J, Lobkovsky AE, Wolf YI, Koonin EV. 2013. Evolutionary Dynamics of the Prokaryotic Adaptive Immunity System CRISPR-Cas in an Explicit Ecological Context. Journal of Bacteriology. 195:3834–3844.

Iranzo J, Lobkovsky AE, Wolf YI, Koonin EV. 2015. Immunity, suicide or both? Ecological determinants for the combined evolution of anti-pathogen defense systems. BMC Evolutionary Biology. 15:43.

Jiang W, Maniv I, Arain F, Wang Y, Levin BR, Marraffini LA. 2013. Dealing with the Evolutionary Downside of CRISPR Immunity: Bacteria and Beneficial Plasmids. PLoS Genet. 9:e1003844.

Kumar MS, Plotkin JB, Hannenhalli S. 2015. Regulated CRISPR Modules Exploit a Dual Defense Strategy of Restriction and Abortive Infection in a Model of Prokaryote-Phage Coevolution. PLoS computational biology. 11:e1004603.

Kuo CH, Ochman H. 2009. Deletional Bias across the Three Domains of Life. Genome Biology and Evolution. 1:145–152.

Kupczok A, Landan G, Dagan T. 2015. The Contribution of Genetic Recombination to CRISPR Array Evolution. Genome Biology and Evolution. 7:1925–1939.

Levy A, Goren MG, Yosef I, Auster O, Manor M, Amitai G, Edgar R, Qimron U, Sorek R. 2015. CRISPR adaptation biases explain preference for acquisition of foreign DNA. Nature. 520:505–510.

Li R, Fang L, Tan S, Yu M, Li X, He S, Wei Y, Li G, Jiang J, Wu M. 2016. Type I CRISPR-Cas targets endogenous genes and regulates virulence to evade mammalian host immunity. Cell Research. 26:1273–1287.

Makarova KS, Grishin NV, Shabalina SA, Wolf YI, Koonin EV. 2006. A putative RNA-interference-based immune system in prokaryotes: computational analysis of the predicted enzymatic machinery, functional analogies with eukaryotic RNAi, and hypothetical mechanisms of action. Biology Direct. 1:7.

Makarova KS, Haft DH, Barrangou R, et al. (12 co-authors). 2011a. Evolution and classification of the CRISPR–Cas systems. Nature Reviews Microbiology. 9:467–477.

Makarova KS, Wolf YI, Alkhnbashi OS, et al. (21 co-authors). 2015. An updated evolutionary classification of CRISPR-Cas systems. Nature Reviews Microbiology. 13:722–736.

Makarova KS, Wolf YI, Koonin EV. 2013. Comparative genomics of defense systems in archaea and bacteria. Nucleic Acids Research. 41:4360–4377.

Makarova KS, Wolf YI, Snir S, Koonin EV. 2011b. Defense islands in bacterial and archaeal genomes and prediction of novel defense systems. Journal of Bacteriology. 193:6039–6056.

Marraffini LA. 2015. CRISPR-Cas immunity in prokaryotes. Nature. 526:55–61.

Marraffini LA, Sontheimer EJ. 2008. CRISPR interference limits horizontal gene transfer in staphylococci by targeting DNA. Science (New York, N.Y.). 322:1843–1845.

Mira A, Ochman H, Moran NA. 2001. Deletional bias and the evolution of bacterial genomes. Trends in genetics: TIG. 17:589–596.

Mojica FJM, Díez-Villaseñor C, García-Martínez J, Soria E. 2005. Intervening Sequences of Regularly Spaced Prokaryotic Repeats Derive from Foreign Genetic Elements. Journal of Molecular Evolution. 60:174–182.

Nivala J, Shipman SL, Church GM. 2018. Spontaneous crispr loci generation in vivo by non-canonical spacer integration. Nature microbiology. p. 1.

O’Leary NA, Wright MW, Brister JR, et al. (55 co-authors). 2016. Reference sequence (RefSeq) database at NCBI: current status, taxonomic expansion, and functional annotation. Nucleic Acids Research. 44:D733–D745.

Paez-Espino D, Sharon I, Morovic W, Stahl B, Thomas BC, Barrangou R, Banfield JF. 2015. CRISPR Immunity Drives Rapid Phage Genome Evolution in Streptococcus thermophilus. mBio. 6:e00262–15.

Pawluk A, Bondy-Denomy J, Cheung VHW, Maxwell KL, Davidson AR. 2014. A New Group of Phage Anti-CRISPR Genes Inhibits the Type I-E CRISPR-Cas System of Pseudomonas aeruginosa. mBio. 5:e00896–14.

Puigbò P, Makarova KS, Kristensen DM, Wolf YI, Koonin EV. 2017. Reconstruction of the evolution of microbial defense systems. BMC Evolutionary Biology. 17:94.

Pyenson NC, Gayvert K, Varble A, Elemento O, Marraffini LA. 2017. Broad targeting specificity during bacterial type iii crispr-cas immunity constrains viral escape. Cell host & microbe. 22:343–353.

Rauch BJ, Silvis MR, Hultquist JF, Waters CS, McGregor MJ, Krogan NJ, Bondy-Denomy J. 2017. Inhibition of CRISPR-Cas9 with Bacteriophage Proteins. Cell. 168:150–158.e10.

Revell LJ. 2012. phytools: an r package for phylogenetic comparative biology (and other things). Methods in Ecology and Evolution. 3:217–223.

Richter H, Zoephel J, Schermuly J, Maticzka D, Backofen R, Randau L. 2012. Characterization of CRISPR RNA processing in Clostridium thermocellum and Methanococcus maripaludis. Nucleic Acids Research. 40:9887–9896.

Savitskaya E, Lopatina A, Medvedeva S, Kapustin M, Shmakov S, Tikhonov A, Artamonova II, Logacheva M, Severinov K. 2017. Dynamics of escherichia coli type i-e crispr spacers over 42 000 years. Molecular ecology. 26:2019–2026.

Shah SA, Garrett RA. 2011. CRISPR/Cas and Cmr modules, mobility and evolution of adaptive immune systems. Research in Microbiology. 162:27–38.

Silas S, Lucas-Elio P, Jackson SA, Aroca-Crevillén A, Hansen LL, Fineran PC, Fire AZ, Sánchez-Amat A. 2017. Type III CRISPR-Cas systems can provide redundancy to counteract viral escape from type I systems. eLife. 6.

Sorek R, Lawrence CM, Wiedenheft B. 2013. CRISPR-Mediated Adaptive Immune Systems in Bacteria and Archaea. Annual Review of Biochemistry. 82:237–266.

Staals RHJ, Jackson SA, Biswas A, Brouns SJJ, Brown CM, Fineran PC. 2016. Interference-driven spacer acquisition is dominant over naive and primed adaptation in a native CRISPR-Cas system. Nature Communications. 7:12853.

Stachler AE, Marchfelder A. 2016. Gene Repression in Haloarchaea using the CRISPR (clustered regularly interspaced short palindromic repeats) - Cas I-B system. Journal of Biological Chemistry. p.jbc.M116.724062.

Stachler AE, Turgeman-Grott I, Shtifman-Segal E, Allers T, Marchfelder A, Gophna U. 2017. High tolerance to self-targeting of the genome by the endogenous CRISPR-Cas system in an archaeon. Nucleic Acids Research..

Stern A, Keren L, Wurtzel O, Amitai G, Sorek R. 2010. Self-targeting by CRISPR: gene regulation or autoimmunity? Trends in Genetics. 26:335–340.

Swarts DC, Mosterd C, van Passel MWJ, Brouns SJJ. 2012. CRISPR Interference Directs Strand Specific Spacer Acquisition. PLoS ONE. 7:e35888.

Tatusova T, DiCuccio M, Badretdin A, Chetvernin V, Nawrocki EP, Zaslavsky L, Lomsadze A, Pruitt KD, Borodovsky M, Ostell J. 2016. NCBI prokaryotic genome annotation pipeline. Nucleic Acids Research. 44:6614–6624.

Touchon M, Charpentier S, Clermont O, Rocha EPC, Denamur E, Branger C. 2011. CRISPR Distribution within the Escherichia coli Species Is Not Suggestive of Immunity-Associated Diversifying Selection. Journal of Bacteriology. 193:2460–2467.

Touchon M, Rocha EPC. 2010. The Small, Slow and Specialized CRISPR and Anti-CRISPR of Escherichia and Salmonella. PLoS ONE. 5:e11126.

Tung Ho Ls, Ané C. 2014. A linear-time algorithm for gaussian and non-gaussian trait evolution models. Systematic biology. 63:397–408.

Tyson GW, Banfield JF. 2008. Rapidly evolving CRISPRs implicated in acquired resistance of microorganisms to viruses. Environmental Microbiology. 10:200–207.

Wei Y, Terns RM, Terns MP. 2015. Cas9 function and host genome sampling in Type II-A CRISPR–Cas adaptation. Genes & Development. 29:356–361.

Weinberger AD, Sun CL, Plucinśki MM, Denef VJ, Thomas BC, Horvath P, Barrangou R, Gilmore MS, Getz WM, Banfield JF. 2012a. Persisting Viral Sequences Shape Microbial CRISPR-based Immunity. PLoS Comput Biol. 8:e1002475.

Weinberger AD, Wolf YI, Lobkovsky AE, Gilmore MS, Koonin EV. 2012b. Viral Diversity Threshold for Adaptive Immunity in Prokaryotes. mBio. 3:e00456–12.

Weissman JL, Holmes R, Barrangou R, Moineau S, Fagan WF, Levin B, Johnson PL. 2018. Immune loss as a driver of coexistence during host-phage coevolution. The ISME journal..

Westra ER, van Houte S, Oyesiku-Blakemore S, Makin B, Broniewski JM, Best A, Bondy-Denomy J, Davidson A, Boots M, Buckling A. 2015. Parasite Exposure Drives Selective Evolution of Constitutive versus Inducible Defense. Current Biology. 25:1043–1049.

Yarza P, Richter M, Peplies J, Euzeby J, Amann R, Schleifer KH, Ludwig W, Glöckner FO, Rosselló-Móra R. 2008. The All-Species Living Tree project: A 16s rRNA-based phylogenetic tree of all sequenced type strains. Systematic and Applied Microbiology. 31:241–250.

Yosef I, Goren MG, Qimron U. 2012. Proteins and DNA elements essential for the CRISPR adaptation process in Escherichia coli. Nucleic Acids Research. p. gks216.

Zeng H, Zhang J, Li C, Xie T, Ling N, Wu Q, Ye Y. 2017. The driving force of prophages and CRISPR-Cas system in the evolution of Cronobacter sakazakii. Scientific Reports. 7:40206.

