## Supplementary Materials for "Selective maintenance of multiple CRISPR arrays across prokaryotes"

### S1 Text Model Analysis

#### Primary Model: Array-Length-Dependent Spacer Loss

Recall the model of spacer dynamics from the main text (Eqs 3-5), where there are $n$ viral species and $C_i$ is the number of spacers targeting viral species $i$

$$\frac{dC_i}{dt} = \begin{cases} \mu_A v_i f_i(t) - \mu_L C_i \sum_{j=1}^n C_j & C_i < 1 \\ p\mu_A v_i f_i(t) - \mu_L C_i \sum_{j=1}^n C_j & C_i \geq 1 \end{cases}. \tag{1}$$

Parameter values can be found in S1 Table.

For the purposes of simplifying our analysis we consider the case where there are two viral species in the system ("background" $B$, and "transient" $T$), one that persists at some background level ($f_B(t) = 1$) and another that returns in sharp bursts at periodic intervals ($f_T(t) \in \{0, 1\}$).

This two-virus situation captures the conflict between the ability to maintain immune memory of recurring infections but also defend against persisting viral enemies. We are interested in the time lag between the return of virus $T$ to the system and the development of host immunity. In the case where memory is maintained during the absence of virus $T$ the lag will be zero, otherwise it will depend on the base spacer uptake rate and the relative densities of the two viral populations.

We examine the time to reacquisition of immunity towards the memory virus using the following procedure (S15 Fig). Note that time is rescaled here to virus return time. Our method is as follows: (1) start the system at its equilibrium state with both viral species present, (2) remove species $T$ for a single unit of time and track the decay of $C_T$, and (3) return $T$ to the system and calculate how long it takes after this return for $C_T$ to exceed one (time to reacquisition of immunity, $t_I$). Let $t_I = 0$ if $C_T$ remains above one despite the absence of $T$ (i.e. no loss of immune memory). In more detail:

1. Find the equilibrium of our system $(\tilde{C}_B, \tilde{C}_T)$ when $T$ is present, assuming that at the equilibrium state $\tilde{C}_B, \tilde{C}_T \geq 1$. We have that

$$\tilde{C}_B = \frac{v_B\sqrt{\mu_A p}}{\sqrt{\mu_L(v_B + v_T)}} \tag{2}$$

and

$$\tilde{C}_T = \frac{v_T\sqrt{\mu_A p}}{\sqrt{\mu_L(v_B + v_T)}}. \tag{3}$$

2. Use this equilibrium as an initial condition for a system where virus $T$ has been removed ($\frac{dC_T}{dt} = -\mu_L C_T\,(C_T + C_B)$) and solve (numerically) for the state of the system at time $t = 1$ assuming that $T$ remains absent ($C_T(1)$).

3. Find the time to reacquisition of immunity ($t_I$ such that $C_T\,(t_I + 1) = 1$) using the solution to the the unprimed system where we assume loss has no effect (since no spacers yet exist to be lost, effect should not be important). Define $C_T^*(t)$ so that

$$C_T^*(t) = \mu_A v_T t + C_T(1). \tag{4}$$

We are interested in $t_I$ where $C_T^*(t_I) = 1$, so that

$$t_I = \frac{1 - C_T(1)}{\mu_A v_T} \tag{5}$$

where $C_T(1)$ is the solution from step (2). This equation only holds if $C_T(1) < 1$ (let $t_I = 0$ otherwise). This time to reacquisition is our measure of fitness, with a lower $t_I$ indicating lower fitness of the host. We define some $T$ such that if $t_I < T$ we say immunity is maintained. Ideally, $T$ should be shorter than the time it takes viruses to cause irreparable damage to an infected bacterium after initial infection.

We can repeat step (3) to find the time of immune acquisition towards a novel viral species ($N$) arriving in the environment. When no spacers towards that species are already contained in the CRISPR array, the time to novel acquisition is

$$t_N = \frac{1}{\mu_A v_N}. \tag{6}$$

For simplicity we let $v_N = v_T$. Note that we assume here that the spacer loss rate does not effect $\frac{dC_N}{dt}$ when $C_N < 1$ in order to simplify our analysis.

#### Alternative Model: Leader-End crRNA Processing

We present a model of a single CRISPR locus targeting $n$ viral species where the change over time of the number of spacers targeting species $i$ is

$$\dot{C}_i = a_i - C_i \sum_j a_j \tag{7}$$

where

$$a_i = \mu v_i f_i(t) g(C_i) \tag{8}$$

so that $\mu$ is the spacer acquisiton rate, $v_i$ is a compound paramter describing the maximal density of viral species $i$ times the adsorption rate, $f_i(t)$ describes the population dynamics of viral species $i$, and the function $g$ incorporates priming so that

$$g(C_i) = \begin{cases} 1 & C_i < \frac{1}{L} \\ p & C_i > \frac{1}{L} \end{cases} \tag{9}$$

where $L$ is the effective length of the locus (number of spacers likely to be transcribed) and $p$ is the degree of priming. Let $\sum_i C_i = 1$ so that $C_i$ represents a proportion of the total "effective" array length ($L$). Observe that in Eq 7 the flow rate into and out of the system match so that the overall number of effective spacers should not change.

When $f_i(t) = 1 \, \forall i$ and we assume the system is primed towards all viruses, then we have equilibrium

$$\tilde{C}_i = \frac{v_i}{\sum_j v_j}. \tag{10}$$

Let's return to our simple two-virus system with background ($B$) and transient ($T$) viral species, concentrating on an interval where both viruses are present ($f_T = 1$)

$$\frac{dC_i}{dt} = \begin{cases} \mu v_i p - p\mu C_i \left(v_i + v_{j\neq i}\right) & C_i, C_{j\neq i} > \frac{1}{L} \\ \mu v_i - \mu C_i \left(v_i + p v_{j\neq i}\right) & C_i < \frac{1}{L}, C_{j\neq i} > \frac{1}{L} \\ \mu v_i p - \mu C_i \left(p v_i + v_{j\neq i}\right) & C_i > \frac{1}{L}, C_{j\neq i} < \frac{1}{L} \\ \mu v_i - \mu C_i \left(v_i + v_{j\neq i}\right) & C_i, C_{j\neq i} < \frac{1}{L} \end{cases} \quad i, j \in \{B, T\}. \tag{11}$$

We derive the equilibrium spacer content, assuming that the system starts from a doubly primed condition so that

$$\tilde{C}_i = \begin{cases} \frac{v_i}{v_i + v_{j\neq i}} & \frac{1}{L} < \frac{v_i}{v_i + v_{j\neq i}} < 1 - \frac{1}{L} \\ \frac{v_i}{v_i + p v_{j\neq i}} & \frac{v_i}{v_i + v_{j\neq i}} < \frac{1}{L} \\ \frac{p v_i}{p v_i + v_{j\neq i}} & \frac{v_i}{v_i + v_{j\neq i}} > 1 - \frac{1}{L} \end{cases} \quad i, j \in \{B, T\}. \tag{12}$$

We now focus on the dynamics of the system after the transient viral species leaves ($f_T = 0$) assuming the system starts from the equilibrium in Eq 12. The decay of spacers targeting the transient viral population will follow

$$\frac{dC_T}{dt} = \begin{cases} -\mu v_B p C_T & C_T < 1 - \frac{1}{L} \\ -\mu v_B C_T & C_T > 1 - \frac{1}{L} \end{cases}. \tag{13}$$

We can solve for $C_T(t)$ with the given initial condition so that

$$C_T(t) = \begin{cases} \left(\frac{v_T}{v_T+v_B}\right) e^{-p\mu v_B t} & \tilde{C}_T, \tilde{C}_B > \frac{1}{L} \\ \left(\frac{v_T}{v_T+pv_B}\right) e^{-p\mu v_B t} & \tilde{C}_T < \frac{1}{L}, \tilde{C}_B > \frac{1}{L} \\ \left(\frac{pv_T}{pv_T+v_B}\right) e^{-\mu v_B t} & t \leq t_B, \tilde{C}_B > \frac{1}{L} \\ \left(\frac{pv_T}{pv_T+v_B}\right) e^{-\mu v_B (t_B + p(t-t_B))} & t > t_B, \tilde{C}_B > \frac{1}{L} \end{cases} \tag{14}$$

where

$$t_B = \frac{-\ln\left(\frac{(pv_T+v_B)(L-1)}{pv_T L}\right)}{\mu v_B}. \tag{15}$$

Let us assume that time is scaled so that the return time of the fluctuation species is $t_R = 1$. Then our goal is to find $C_T(1)$, assess whether this value has dropped below $\frac{1}{L}$ (immune loss), and if so, find the time to immune reacquisition $C_T(t_I - 1) = \frac{1}{L}$. Let us define a new function $C_T^*(t)$ that tracks the re-acquisition of immunity after $T$ returns to the system so that $C_T^*(0) = C_T(1)$ and, assuming no decay of spacers when $C_T^* < \frac{1}{L}$,

$$\frac{1}{L} = C_T^*(t_I) = \mu v_T t_I + C_T(1). \tag{16}$$

Then

$$t_I = \frac{\frac{1}{L} - C_T(1)}{\mu v_T} \tag{17}$$

and we have $C_T(1)$ from Eq 14.

This model gives qualitatively similar results to those found with the primary model (S10 Fig).

### S2 Text Autoimmunity Constrains Unprimed Spacer Acquisition Rates

Empirical evidence suggests that there is no self versus non-self recognition mechanism in the CRISPR systems of *Streptococcus thermophilus*, a popular model system for CRISPR research [1]. Thus any increase in spacer acquisition will also increase the rate of autoimmune targeting. In the absence of viruses or once immunity has been established, we can model the growth of bacteria ($B$) experiencing autoimmune targeting in a chemostat as

$$\dot{B} = B\left(\frac{vR}{z+R} - \alpha - w\right) \tag{18}$$

with resources ($R$)

$$\dot{R} = w(A-R) - \frac{evR}{R+z}B \tag{19}$$

where $\mu_A$ is the spacer acquisition rate and $\alpha = 50\mu_A$ is an estimate of the rate of autoimmunity based of the relative genome sizes of *S. thermophilus* and its lytic phage 2972 [2, 3] (other parameters in S2 Table).

As shown in S16 Fig, there is little to no effect of autoimmunity on the equilibrium density of bacteria for $\mu_A < 10^{-5}$, but after a point around $\mu_A = 10^{-4}$ there is a rapid drop in density. This puts a theoretical cap on the maximum rate of spacer uptake in our system and imposes a severe cost on spacer uptake rates greater than $10^{-4}$ given the parameters used here, based on the *S. thermophilus* system. Other taxa do seem to exhibit some degree of self versus non-self recognition, but still frequently incorporate self-spacers [4, 5, 6], suggesting that our general result holds across taxa though the threshold is likely to be variable. We note that in S16 Fig even a 50% reduction in the rate of autoimmunity only shifts the threshold spacer acquisition rate by a small amount.

### S3 Text Bet Hedging Against Memory Loss

Spacer loss in the CRISPR array most likely occurs via homologous recombination of repeat sequences [7, 8, 9]. Thus the time to immune loss will increase with the number of arrays targeting a particular viral species. Assuming that immunity towards a given virus in a single array has an exponentially distributed lifetime with expected value $\tau$ (i.e., time to loss of all spacers targeting that virus in that array), in the absence of novel acquisitions the expected time to complete immune loss is $\tau \sum_{i=1}^{N} \frac{1}{i}$, where $N$ is the number of arrays that initially target the virus in question. Clearly, the advantage conferred in terms of memory span decreases with each additional array, though this effect is important for the first few added arrays. In fact, it is more appropriate to model the lifetime of individual spacers with an exponential distribution such that the expected time to complete immune loss is $\tau_s \sum_{i=1}^{n} \frac{1}{i}$, where $n$ is the total number of spacers in all arrays and $\tau_s$ the expected lifetime of each spacer. Thus the relative advantage of multiple arrays is further reduced in the case where each array can have multiple spacers targeting the same virus, assuming that spacer loss rates are similar across arrays (appropriate in the case of identical arrays near some equilibrium length).

If spacers vary in their effectiveness in attacking a viral target then we would expect this to increase the relative payoff of a bet-hedging strategy since it will essentially reduce the number of effective spacers in any given array. There is evidence that spacers vary in their targeting efficiency [10] in some systems. Nevertheless, if a system experiences priming then it is extremely likely that a single array would have many spacers towards the same target, making a bet hedging strategy less likely.

### References


[1] Wei Y, Terns RM, Terns MP. Cas9 function and host genome sampling in Type II-A CRISPR–Cas adaptation. Genes & Development. 2015;29(4):356–361. doi:10.1101/gad.257550.114.

[2] Bolotin A, Quinquis B, Sorokin A, Ehrlich SD. Clustered regularly interspaced short palindrome repeats (CRISPRs) have spacers of extrachromosomal origin. Microbiology. 2005;151(8):2551–2561. doi:10.1099/mic.0.28048-0.

[3] Lévesque C, Duplessis M, Labonté J, Labrie S, Fremaux C, Tremblay D, et al. Genomic Organization and Molecular Analysis of Virulent Bacteriophage 2972 Infecting an Exopolysaccharide-Producing Streptococcus thermophilus Strain. Applied and Environmental Microbiology. 2005;71(7):4057–4068. doi:10.1128/AEM.71.7.4057-4068.2005.

[4] Stern A, Keren L, Wurtzel O, Amitai G, Sorek R. Self-targeting by CRISPR: gene regulation or autoimmunity? Trends in Genetics. 2010;26(8):335–340. doi:10.1016/j.tig.2010.05.008.

[5] Yosef I, Goren MG, Qimron U. Proteins and DNA elements essential for the CRISPR adaptation process in Escherichia coli. Nucleic Acids Research. 2012; p. gks216. doi:10.1093/nar/gks216.

[6] Levy A, Goren MG, Yosef I, Auster O, Manor M, Amitai G, et al. CRISPR adaptation biases explain preference for acquisition of foreign DNA. Nature. 2015;520(7548):505–510. doi:10.1038/nature14302.

[7] Garrett RA, Shah SA, Vestergaard G, Deng L, Gudbergsdottir S, Kenchappa CS, et al. CRISPR-based immune systems of the Sulfolobales: complexity and diversity. Biochemical Society Transactions. 2011;39(1):51–57. doi:10.1042/BST0390051.

[8] Gudbergsdottir S, Deng L, Chen Z, Jensen JVK, Jensen LR, She Q, et al. Dynamic properties of the Sulfolobus CRISPR/Cas and CRISPR/Cmr systems when challenged with vector-borne viral and plasmid genes and protospacers. Molecular Microbiology. 2011;79(1):35–49. doi:10.1111/j.1365-2958.2010.07452.x.

[9] Weinberger AD, Sun CL, Pluciński MM, Denef VJ, Thomas BC, Horvath P, et al. Persisting Viral Sequences Shape Microbial CRISPR-based Immunity. PLoS Comput Biol. 2012;8(4):e1002475. doi:10.1371/journal.pcbi.1002475.

[10] Xue C, Seetharam AS, Musharova O, Severinov K, J Brouns SJ, Severin AJ, et al. CRISPR interference and priming varies with individual spacer sequences. Nucleic Acids Research. 2015;43(22):10831–10847. doi:10.1093/nar/gkv1259.

[11] Van Orden MJ, Klein P, Babu K, Najar FZ, Rajan R. Conserved DNA motifs in the type II-A CRISPR leader region. PeerJ. 2017;5:e3161. doi:10.7717/peerj.3161.

| Symbol | Definition | Value |
|---|---|---|
| $\mu_A$ | Unprimed Spacer Acquisition Rate | varied Fig 2a and S9 Fig |
| $\mu_L$ | Per-Spacer Loss Rate | $10^{-2}$ |
| $v_T$ | Density Virus $T\times$Adsorption | $10^2$ |
| $v_B$ | Density Virus $B\times$Adsorption | $10^2$, varied in Fig 2a |
| $p$ | Priming Factor | $10^4$, varied in S9 Fig |

S1 Table: Definitions of relevant variables and parameters for CRISPR array model.

| Symbol | Definition | Value |
|---|---|---|
| $e$ | Resource Consumption Rate of Growing Bacteria | $5 \times 10^{-7} \mu$g/mL |
| $v$ | Maximum Bacterial Growth Rate | 1.4 divisions/hr |
| $z$ | Resource Concentration for Half-Maximal Growth | 1 $\mu$g/mL |
| $w$ | Flow Rate | 0.3 mL/hr |
| $A$ | Resource Pool | 350 $\mu$g/mL |

S2 Table: Definitions of relevant variables and parameters for autoimmunity model.

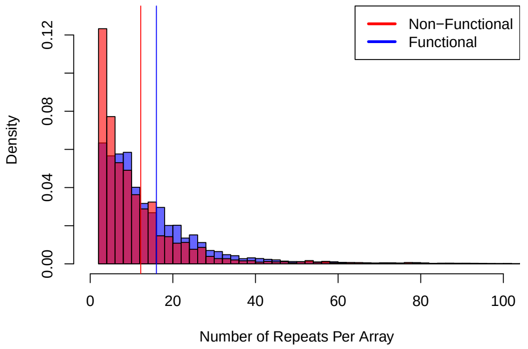


(a)

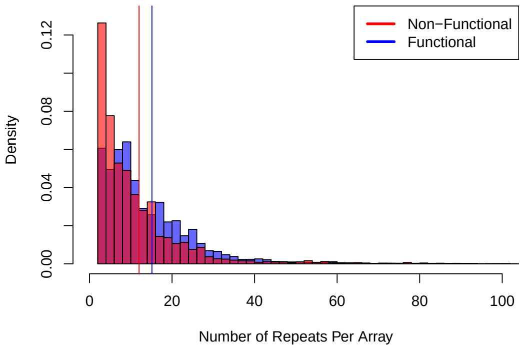


(b)

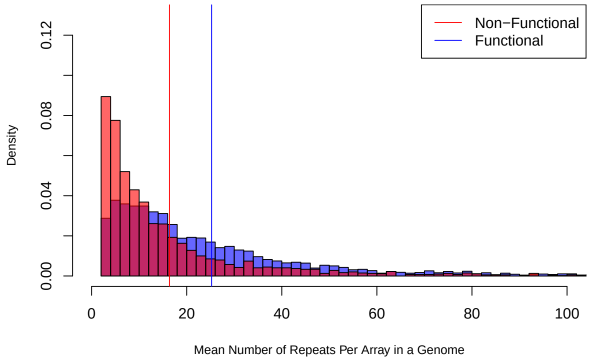


(c)

S1 Fig: **Arrays in functional genomes are longer on average than arrays in non-functional genomes (t-test, $p < 2 \times 10^{-16}$).** (a) Full dataset. (b) Data with a single set of *cas* genes or less shown. (c) Functional genomes tend to have more repeats on average in their CRISPR arrays than non-functional genomes (array length first averaged over arrays in each genome). In blue is the distribution of mean array length in functional genomes. In red (overlaid) is the distribution of mean array length in non-functional genomes.

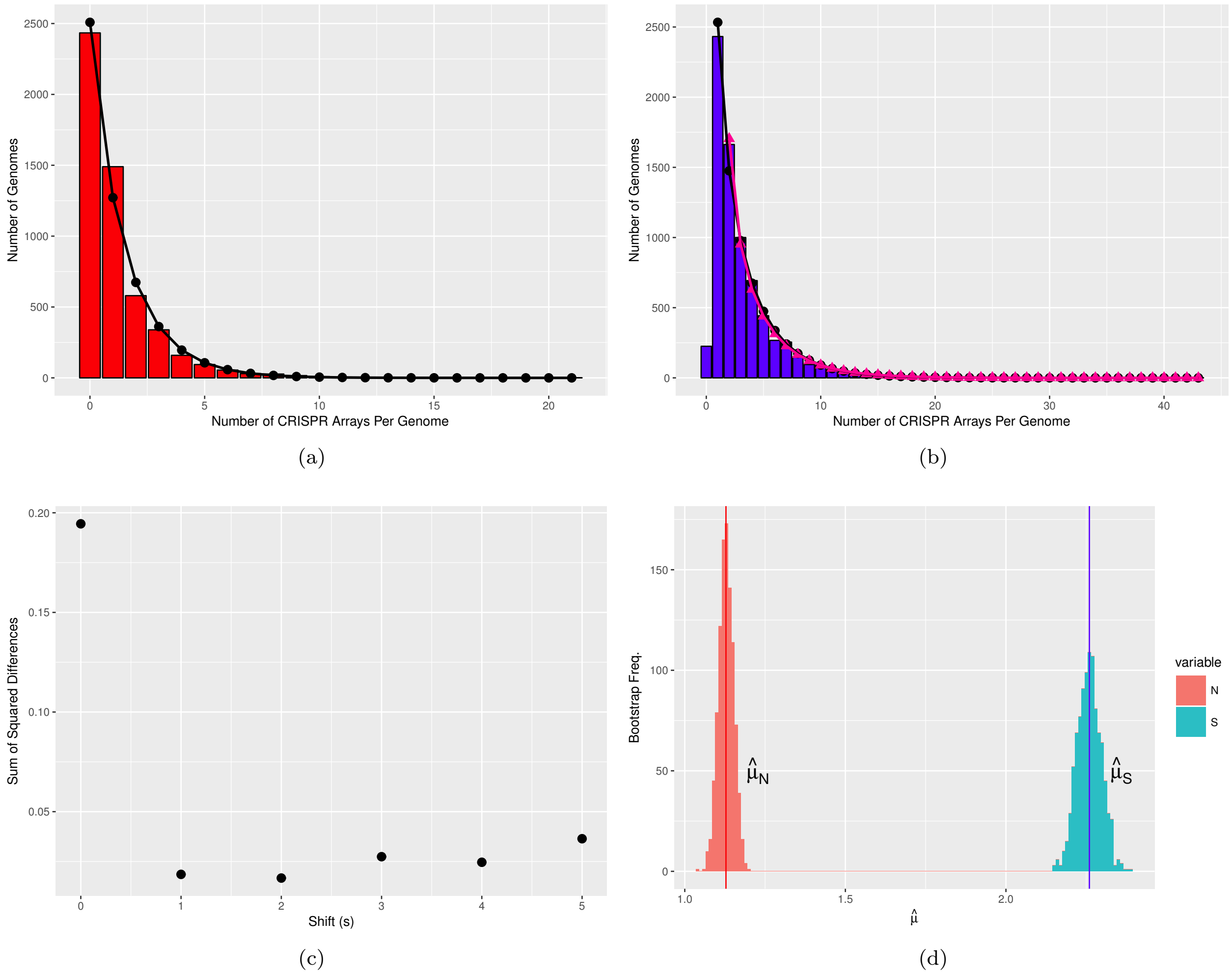


S2 Fig: **Dataset with subsampled genomes of overrepresented taxa.** (a-b) Distribution of number of arrays per genome in (a) non-functional genomes and (b) functional genomes. In (a) the black circles show the negative binomial fit to the distribution of arrays in non-functional genomes. In (b) black circles indicate the negative binomial fit to the single-shifted distribution ($s = 1$) and pink triangles to the double-shifted distribution ($s = 2$). (c) The optimal shift is where the differences between the two distributions is minimized. (d) The bootstrapped distributions of the parameter estimates of $\hat{\mu}_S$ and $\hat{\mu}_N$ show no overlap with $n = 1000$ samples drawn.

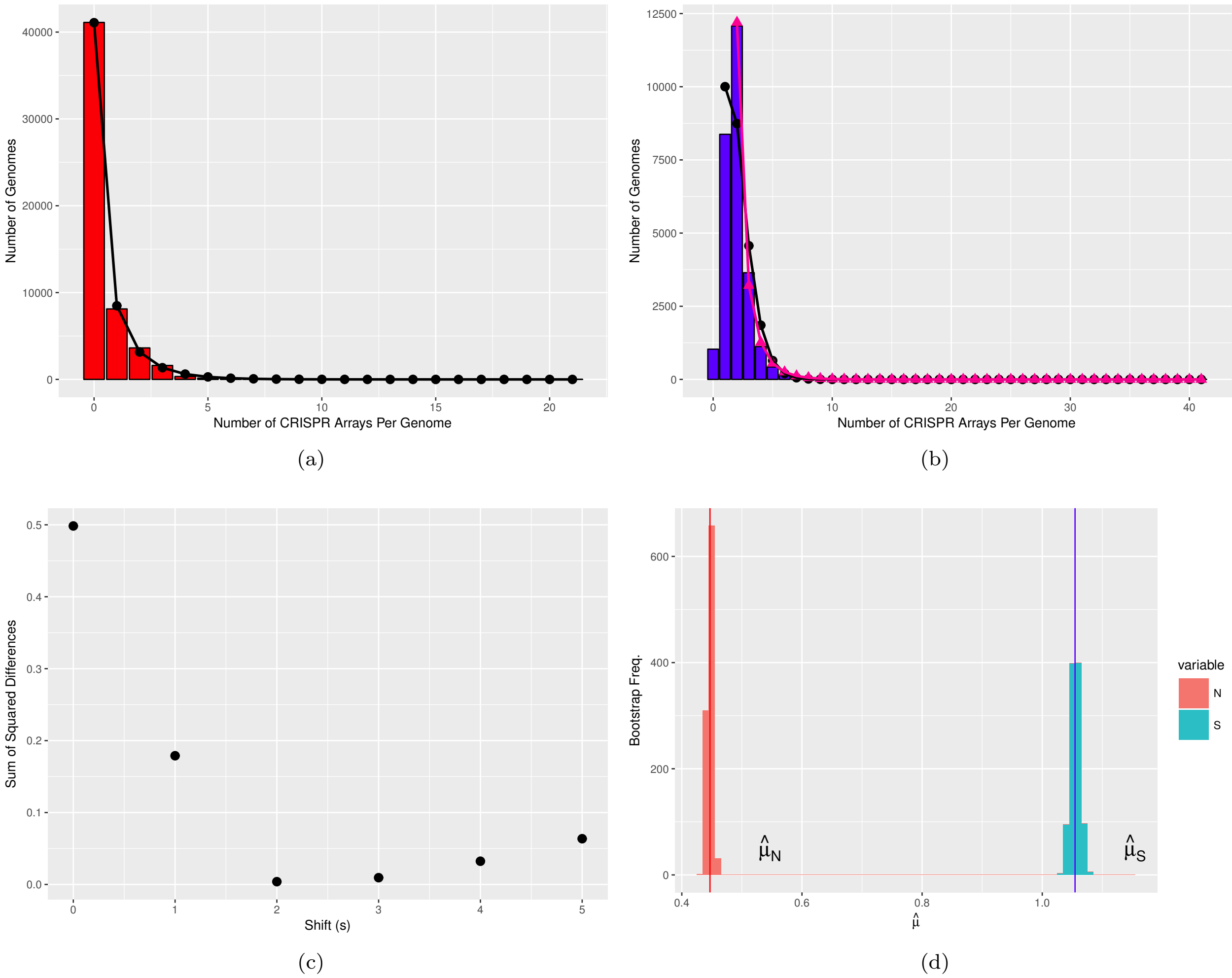


S3 Fig: **Dataset restricted to genomes with a single set of or no *cas* genes (one copy or less each of *cas1*, *cas2*, and a *cas* targeting gene).** (a-b) Distribution of number of arrays per genome in (a) non-functional genomes and (b) functional genomes. In (a) the black circles show the negative binomial fit to the distribution of arrays in non-functional genomes. In (b) black circles indicate the negative binomial fit to the single-shifted distribution ($s = 1$) and pink triangles to the double-shifted distribution ($s = 2$). (c) The optimal shift is where the differences between the two distributions is minimized. (d) The bootstrapped distributions of the parameter estimates of $\hat{\mu}_S$ and $\hat{\mu}_N$ show no overlap with $n = 1000$ samples drawn.

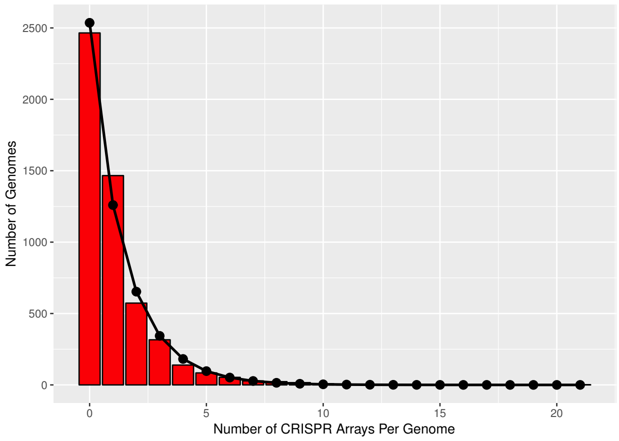


(a)

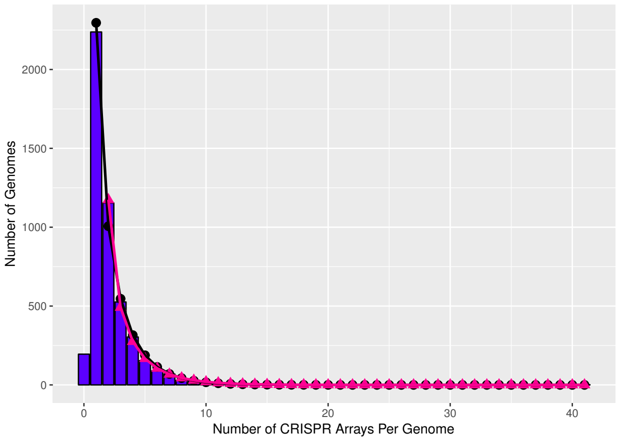


(b)

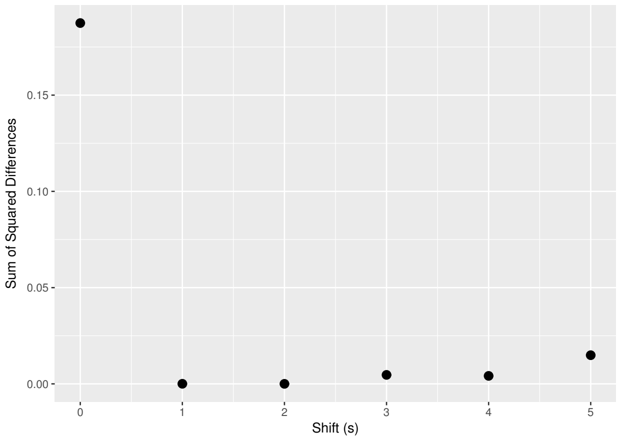


(c)

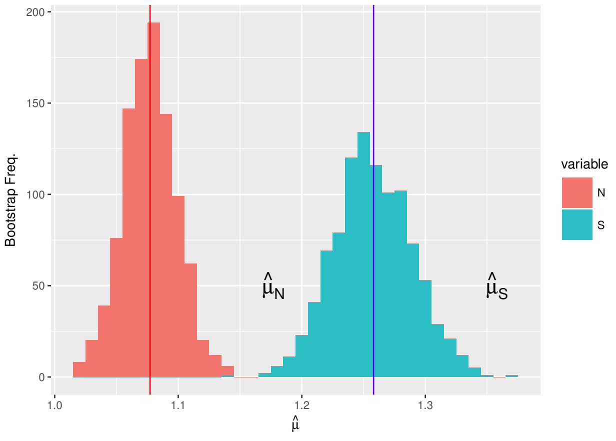


(d)

S4 Fig: **Dataset restricted to genomes with a single set of or no *cas* genes (one copy or less each of *cas1*, *cas2*, and a *cas* targeting gene) with subsampled genomes of overrepresented taxa.** (a-b) Distribution of number of arrays per genome in (a) non-functional genomes and (b) functional genomes. In (a) the black circles show the negative binomial fit to the distribution of arrays in non-functional genomes. In (b) black circles indicate the negative binomial fit to the single-shifted distribution ($s = 1$) and pink triangles to the double-shifted distribution ($s = 2$). (c) The optimal shift is where the differences between the two distributions is minimized. (d) The bootstrapped distributions of the parameter estimates of $\hat{\mu}_S$ and $\hat{\mu}_N$ show no overlap with $n = 1000$ samples drawn.

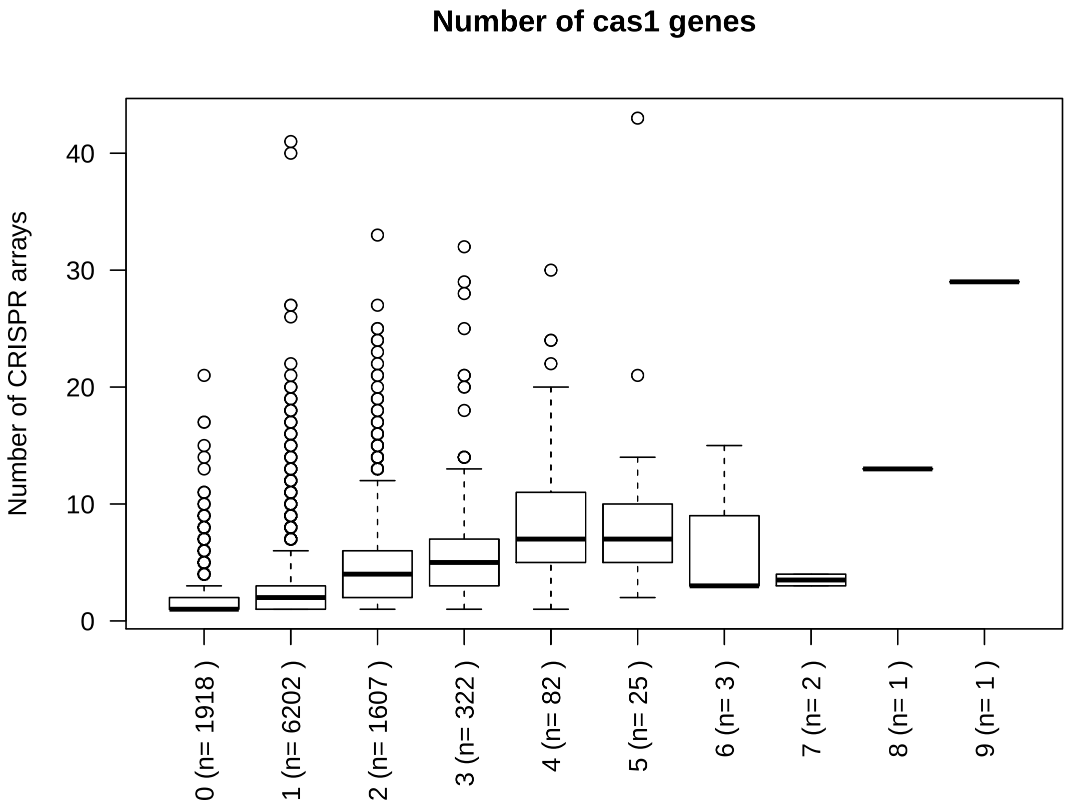


S5 Fig: **Relationship between the number of *cas1* genes and the number of CRISPR arrays in a genome.**

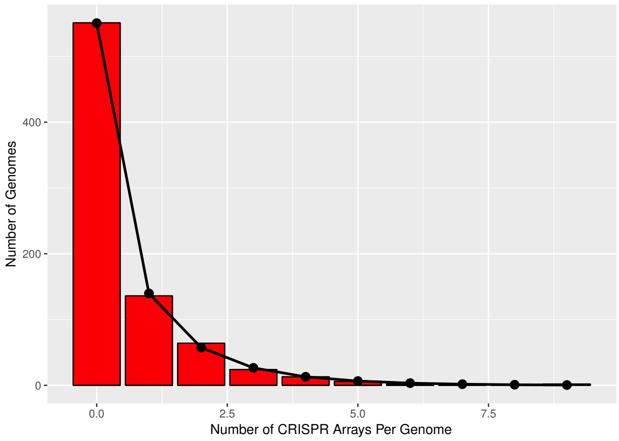


(a)

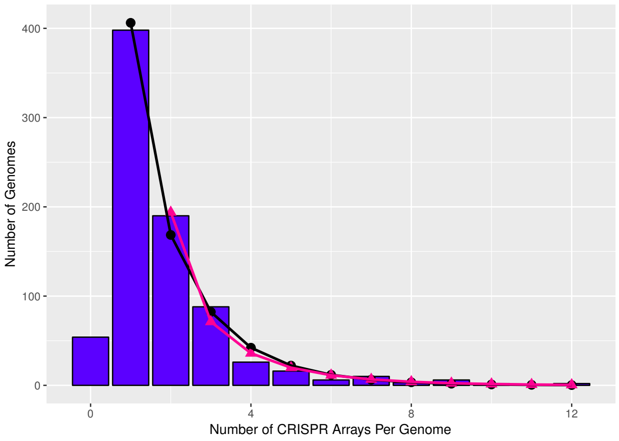


(b)

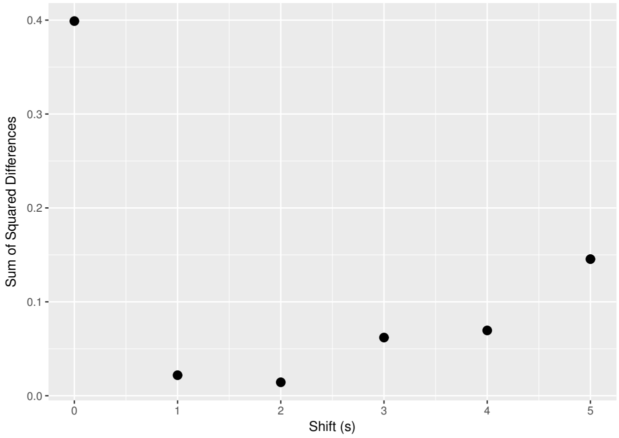


(c)

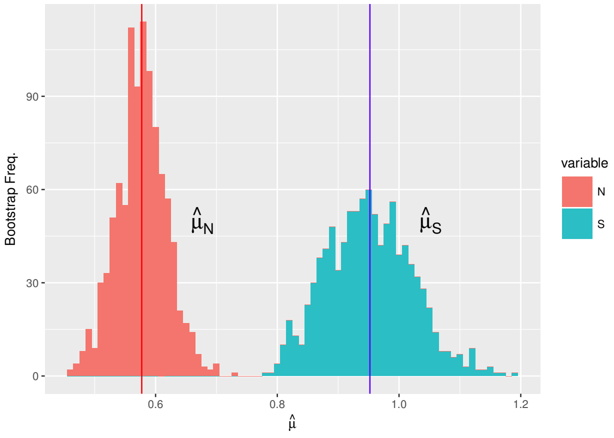


(d)

S6 Fig: **Dataset restricted to genomes with a single set of or no *cas* genes with subsampled genomes of overrepresented taxa, subsetted such that each species is represented by an equal number of genomes with and without *cas* genes.** (a-b) Distribution of number of arrays per genome in (a) non-functional genomes and (b) functional genomes. In (a) the black circles show the negative binomial fit to the distribution of arrays in non-functional genomes. In (b) black circles indicate the negative binomial fit to the single-shifted distribution ($s = 1$) and pink triangles to the double-shifted distribution ($s = 2$). (c) The optimal shift is where the differences between the two distributions is minimized. (d) The bootstrapped distributions of the parameter estimates of $\hat{\mu}_S$ and $\hat{\mu}_N$ show no overlap with $n = 1000$ samples drawn.

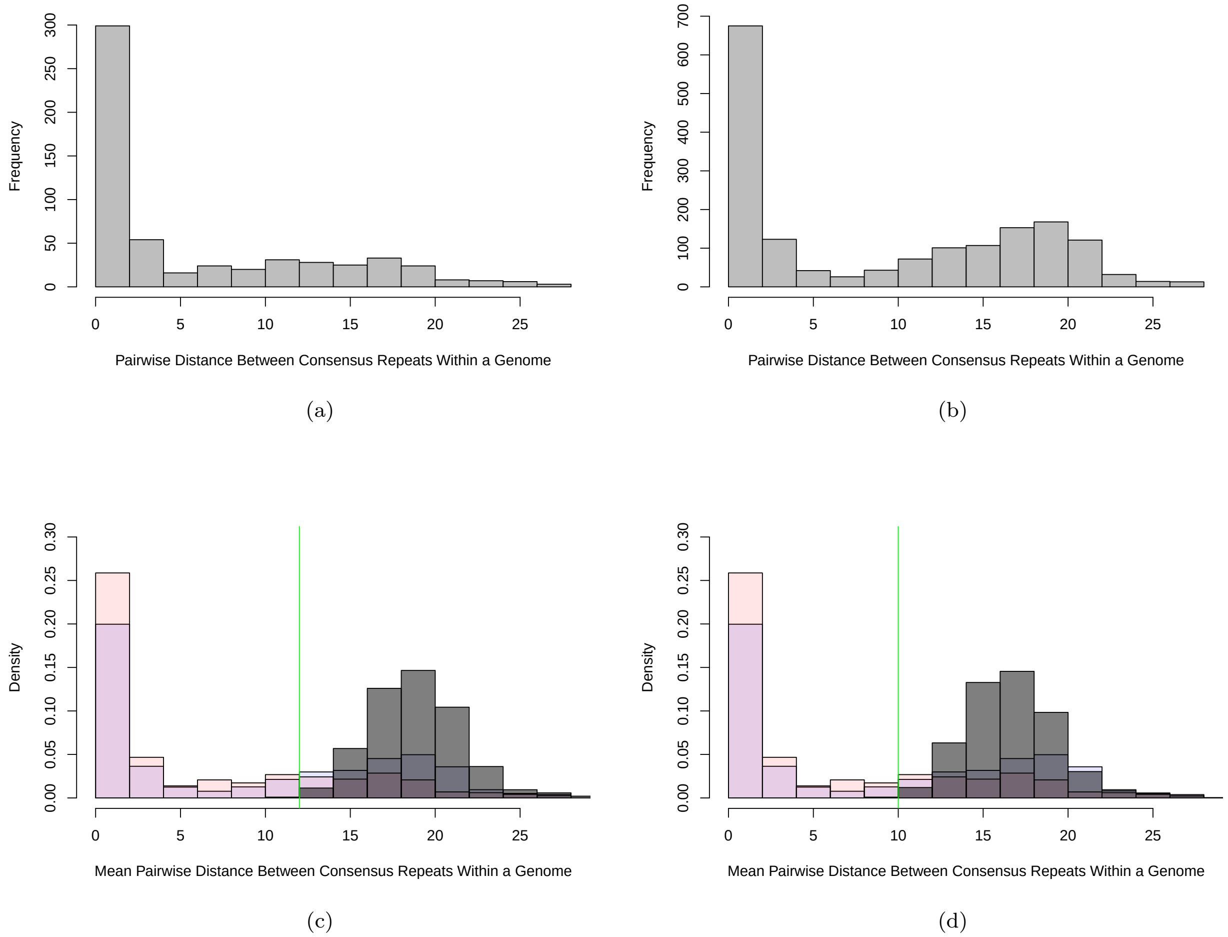


S7 Fig: **Pairwise distance between consensus repeats from arrays within a genome (only genomes with two arrays shown).** Distance calculated as Levenshtein Distance between each pair divided by the length of the longest repeat in the pair. (a) Non-functional genomes. (b) Functional genomes. genomes. The functional genomes are enriched with dissimilar arrays ($\chi^2 = 26.406$, $df = 1$, $p = 2.766 \times 10^{-7}$, dissimilarity cutoff at 3 based on median across all two-array genomes). (c) The distributions of pairwise differences between arrays in functional genomes (blue, a) and non-functional genomes (red, b) are overlaid alongside a distribution of distances between random sequences with lengths drawn from the empirical distribution of repeat lengths in the full dataset (gray). The green line indicates the bottom 0.1% of this random (gray) distribution, which can be used as an alternative similarity cutoff ($\chi^2 = 54.653$, $df = 1$, $p = 1.505 \times 10^{-13}$). (d) Same as (c) but with 4 bases held constant across all repeats to simulate some degree of universal sequence conservation at one end of the repeat as observed among type II-A CRISPR systems [11] ($\chi^2 = 64.168$, $df = 1$, $p = 1.142 \times 10^{-15}$). In (c,d) the simulated distribution takes into account the overall frequency of each base across repeat sequences. Histograms drawn using default settings of hist() function in R (right-closed/left-open intervals, except for the first interval which includes the lower bound, i.e. zero).

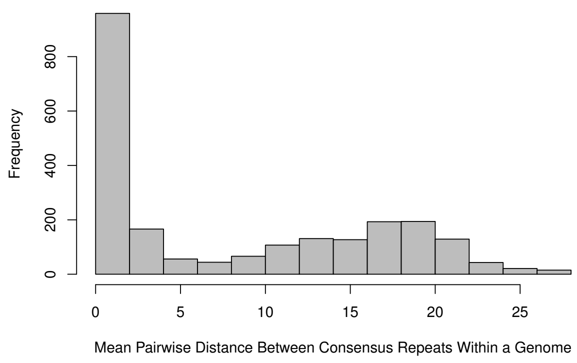


(a)

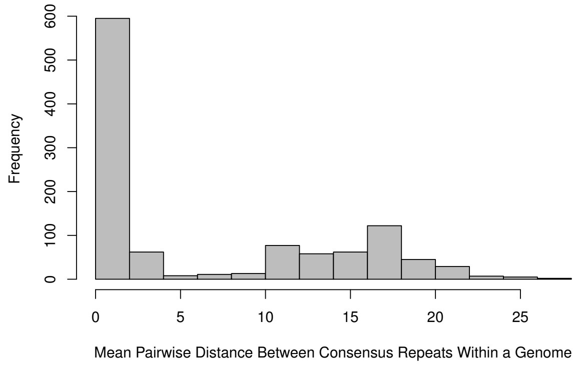


(b)

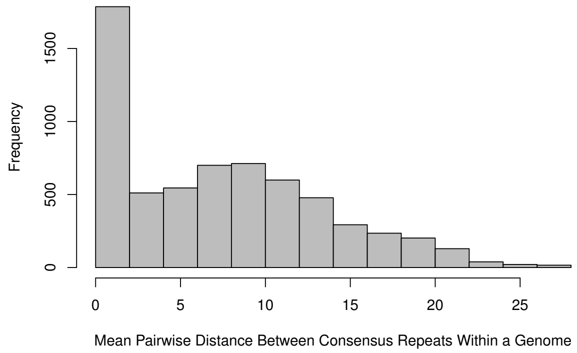


(c)

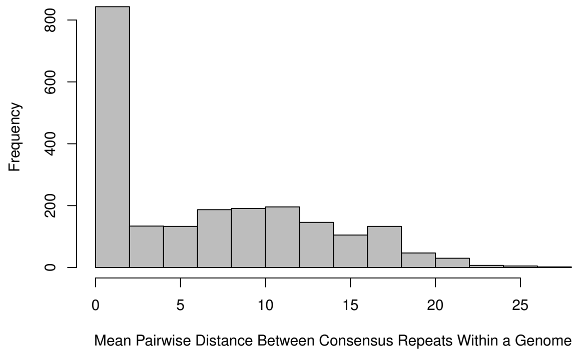


(d)

S8 Fig: **Consensus repeat diversity among arrays in a genome.** (a) Pairwise distance between consensus repeats from arrays within a genome (only genomes with exactly two arrays shown). Distance calculated as Levenshtein Distance between each pair. Histogram drawn using default settings of hist() function in R (right-closed/left-open intervals, except for the first interval which includes the lower bound, i.e. zero). (b) The same as (a) for the CRISPR Database dataset. (c,d) Mean pairwise distance between consensus repeats from all arrays in a genome, including genomes with more than two arrays, for (c) the CRISPRDetect and (d) the CRISPR Database datasets.

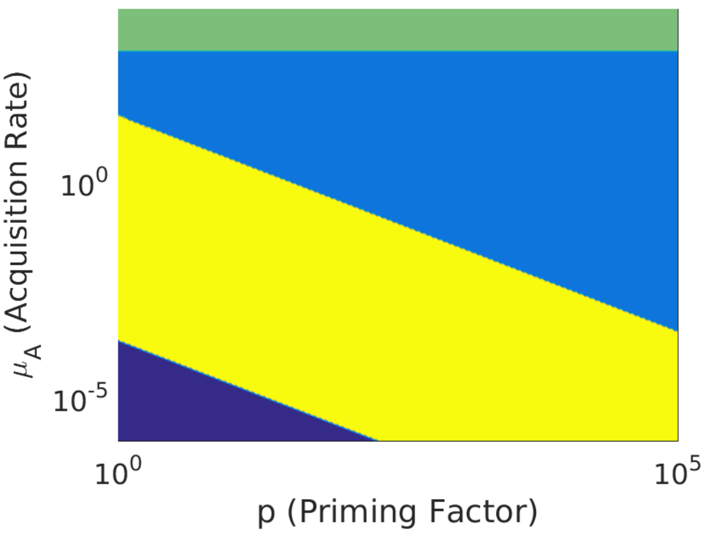


S9 Fig: **Priming increases the region of memory washout and thus deepens the memory span versus acquisition rate tradeoff.** Phase diagram of the behavior of our CRISPR array model with two viral species, a constant “background” population and a “transient” population that leaves and returns to the system at some fixed interval (S1 Text, S13 Fig). The yellow region indicates that immunity towards both viral species was maintained. The green region indicates where immune memory towards the transient viral species was lost, but reacquired almost immediately upon viral reintroduction. The light blue region indicates that only immunity towards the background species was maintained (i.e., immune memory towards the transient viral species was rapidly lost but not rapidly reacquired). Dark blue indicates where equilibrium spacer content towards one or both species did not exceed one despite both species being present in the system. The parameter $p$ is the priming factor by which acquisition rate is increased when spacers towards a given target already exist in the array.

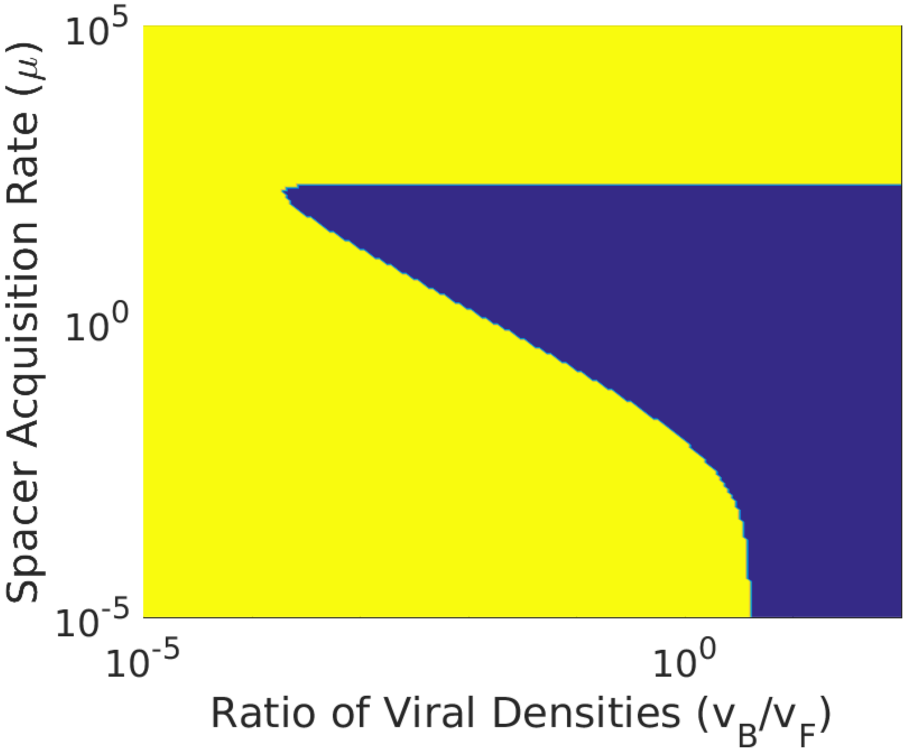


S10 Fig: **Alternative model results with array length cap agree qualitatively with those of the primary model ($p = 1$ (no priming), $L = 5$, and $v_T = 100$).** Blue signifies memory washout ($t_I \geq 10^{-5}$) and yellow signifies immune maintenance ($t_I < 10^{-5}$).

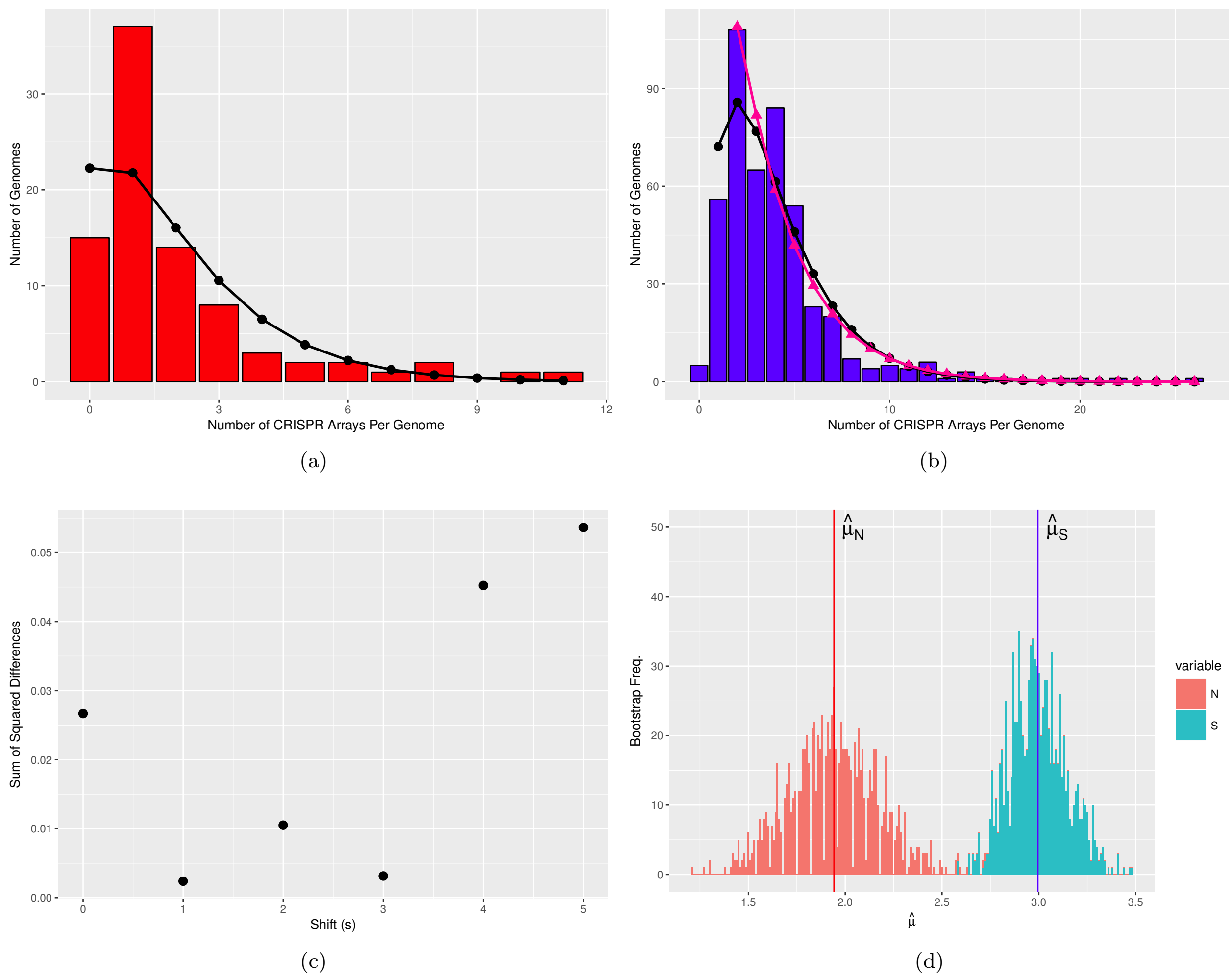


S11 Fig: **Signature of multi-array selection in archaeal genomes.** (a-b) Distribution of number of arrays per genome in (a) non-functional genomes and (b) functional genomes. In (a) the black circles show the negative binomial fit to the distribution of arrays in non-functional genomes. In (b) black circles indicate the negative binomial fit to the single-shifted distribution ($s = 1$) and pink triangles to the double-shifted distribution ($s = 2$). (c) The optimal shift is where the differences between the two distributions is minimized. (d) The bootstrapped distributions of the parameter estimates of $\hat{\mu}_S$ and $\hat{\mu}_N$ show significant overlap with $n = 1000$ samples drawn.

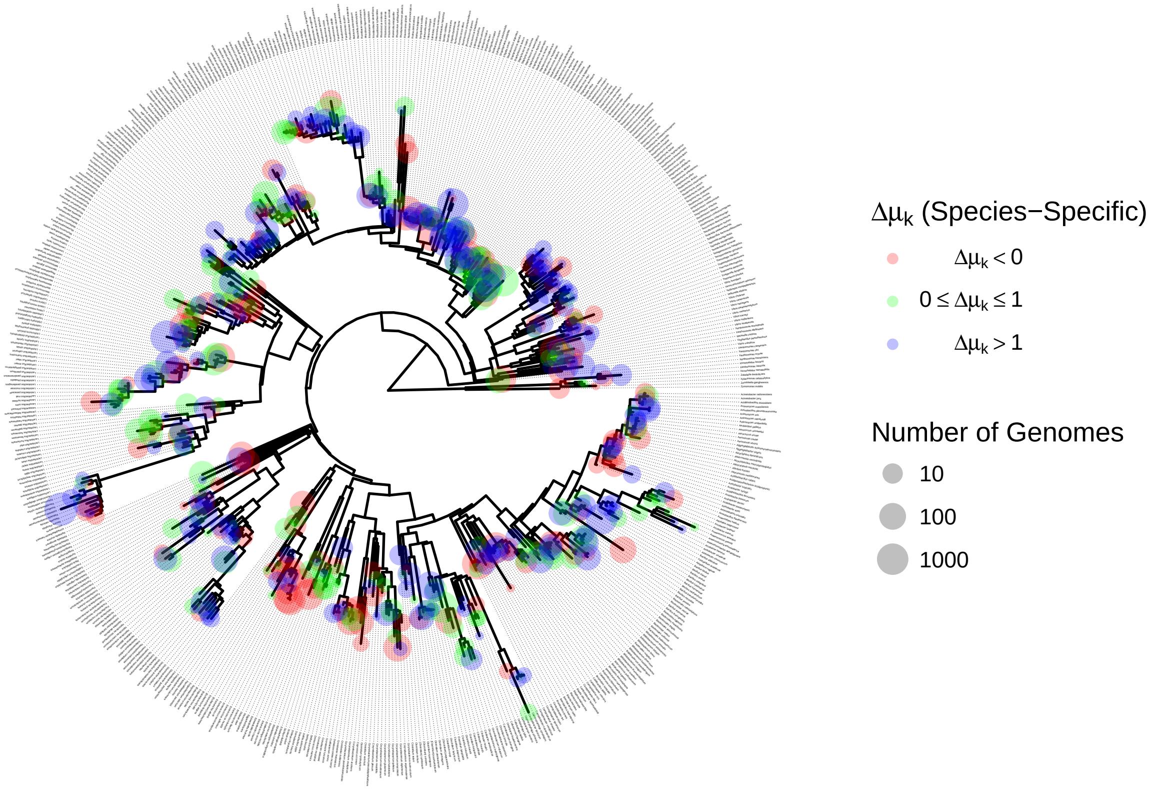


S12 Fig: **Species-specific $\Delta\mu_k$ values mapped onto the SILVA Living Tree 16s rRNA Tree.** In red are species experiencing apparent selection against having a functional CRISPR array. In blue are species showing a strong signature of selection for multiple arrays. Number of genomes is represented by tip size, and is a rough indicator of power.

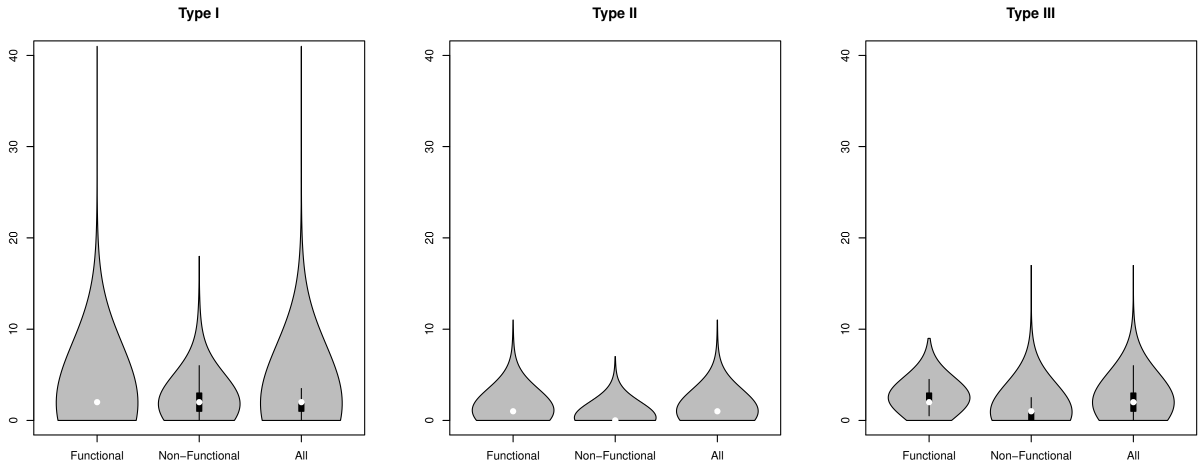


S13 Fig: **Violin plots of array counts associated with genomes carrying a particular type of *cas* targeting machinery.** System type was determined by the type of *cas* targeting gene found on the genome (genomes with no signature targeting genes or multiple types are excluded)

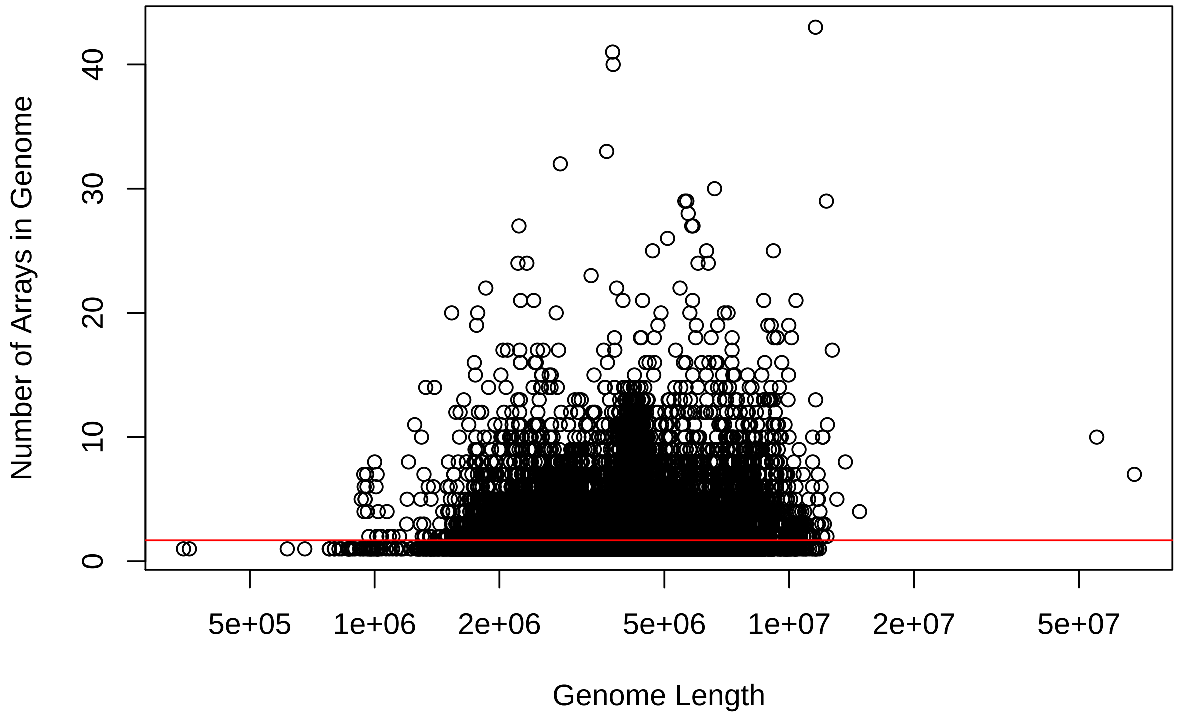


S14 Fig: **Genome length does not explain extreme array counts in some genomes.** While there is a weak positive association between genome length and the number of arrays in a genome ($m = 1.457 \times 10^{-7}$, $R^2 = 0.01143$, $p = 2.2 \times 10^{-16}$), we do not observe an obvious relationship between extreme array counts and genome length.

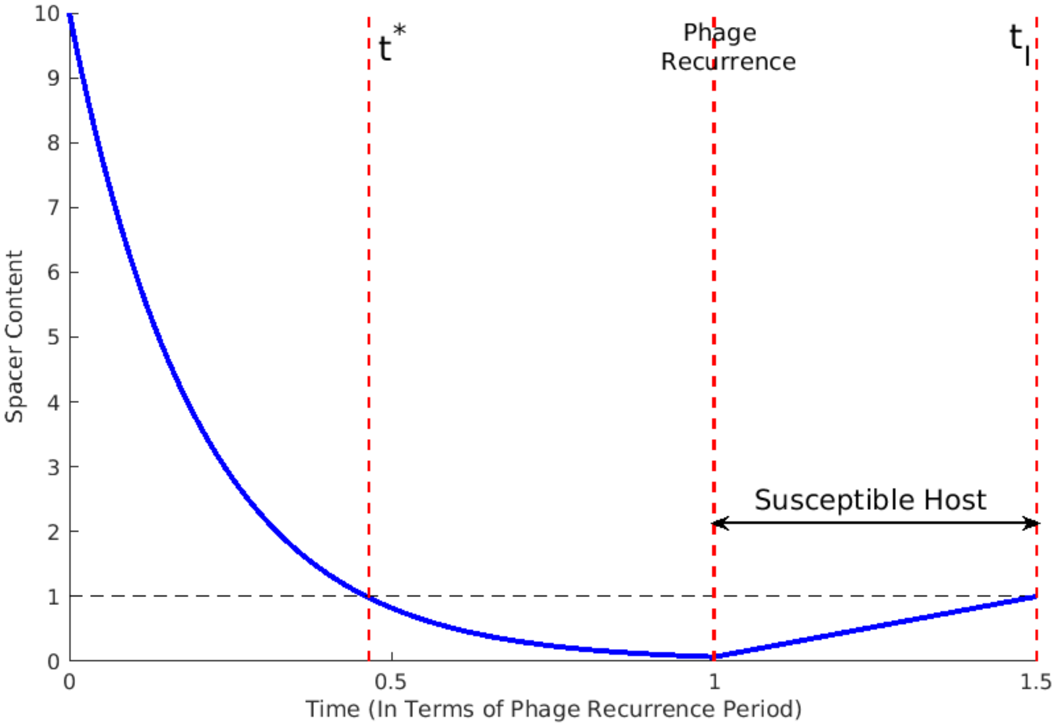


S15 Fig: **Outline of model analysis.** Hypothetical time-course of $C_T$ during the departure and return of virus $T$ from the system. Virus $T$ leaves at time $t = 0$ and returns at time $t = 1$. $C_T$ exceeds a value of one after viral reintroduction at time $1 + t_I$.

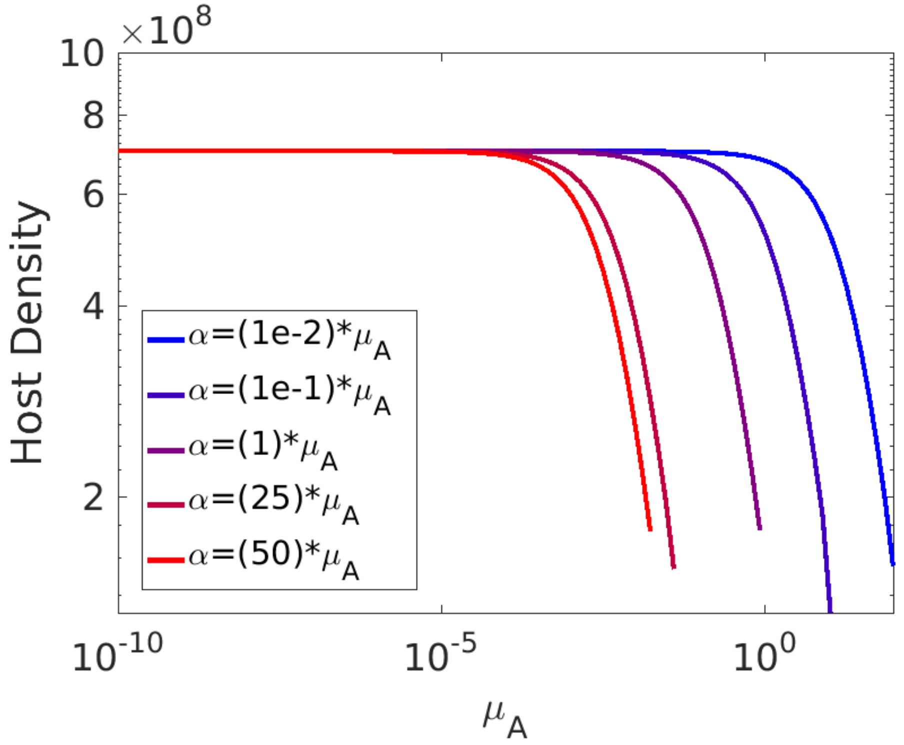


S16 Fig: **Equilibrium host density values from autoimmunity model (S2 Text) over varying spacer acquisition rates.** Curves end because for extremely high $\alpha$ the equilibrium no longer exists if we restrict both host density and resource concentration to be non-negative.
